# The evolutionary dynamics and fitness landscape of clonal haematopoiesis

**DOI:** 10.1101/569566

**Authors:** Caroline J. Watson, Alana Papula, Yeuk P. G. Poon, Wing H. Wong, Andrew L. Young, Todd E. Druley, Daniel S. Fisher, Jamie R. Blundell

**Affiliations:** Department of Oncology, University of Cambridge, United Kingdom; Early Detection Programme, CRUK Cambridge Cancer Centre, University of Cambridge, United Kingdom; Department of Applied Physics, Stanford University, California, USA; Department of Pediatrics, Division of Hematology and Oncology, Washington University School of Medicine, St. Louis, USA

**Keywords:** clonal haematopoiesis, haematopoietic stem cells, evolution, population genetics, DNMT3A, TET2, spliceosome, TP53, acute myeloid leukaemia

## Abstract

Somatic mutations acquired in healthy tissues as we age are major determinants of cancer risk. Whether variants confer a fitness advantage or rise to detectable frequencies by chance, however, remains largely unknown. Here, by combining blood sequencing data from ∼50,000 individuals, we reveal how mutation, genetic drift and fitness differences combine to shape the genetic diversity of healthy blood (‘clonal haematopoiesis’). By analysing the spectrum of variant allele frequencies we quantify fitness advantages for key pathogenic variants and genes and provide bounds on the number of haematopoietic stem cells. Positive selection, not drift, is the major force shaping clonal haematopoiesis. The remarkably wide variation in variant allele frequencies observed across individuals is driven by chance differences in the timing of mutation acquisition combined with differences in the cell-intrinsic fitness effect of variants. Contrary to the widely held view that clonal haematopoiesis is driven by ageing-related alterations in the stem cell niche, the data are consistent with the age dependence being driven simply by continuing risk of mutations and subsequent clonal expansions that lead to increased detectability at older ages.

---

As we age, physiologically healthy tissues such as skin ^1, 2^, colon ^3, 4^, oesophagus ^5, 6^ and blood ^7–18^ acquire mutations in cancer-associated genes. In blood this phenomenon, termed clonal haematopoiesis (CH), increases in prevalence with age ^7–18^, becoming almost ubiquitous in those over the age of 65^10, 15^. The majority of CH mutations are thought to arise in haematopoietic stem cells (HSCs) ^10, 19^ and typically fall within the genes DNMT3A, TET2, ASXL1, JAK2, TP53 and spliceosome genes, although chromosomal alterations are also observed ^17^. Because CH is associated with an increased risk of blood cancers ^7,8,19^, and the genes affected are commonly mutated in pre-leukaemic stem cells ^20–24^, CH has emerged as an important pre-cancerous state, for which a quantitative understanding would accelerate risk stratification and improve our understanding of normal haematopoiesis.

The risk of progressing to a blood cancer depends on the gene in which a variant falls ^14, 18^. However, our ability to risk stratify specific variants remains crude. Should all DNMT3A variants be considered low risk and all spliceosome variants considered equally high risk? If variants confer a fitness advantage to HSCs they are more likely to expand over time, and higher variant allele frequencies (VAFs) are strong predictors of AML development ^14, 18^. It stands to reason there-fore, that by analysing the spectrum of VAFs, one might be able infer the fitness advantage conferred by variants, even from a static ‘snapshot’. This would enable us to generate a comprehensive map between specific variants and their fitness consequences, allowing risk to be stratified with greater resolution.

A major challenge to using VAFs to risk stratify variants is that the spectrum of VAFs, even for a single variant, is remarkably broad, varying by over three orders-of-magnitude across individuals. Whether these differences in VAFs are a result of cell-intrinsic fitness advantages ^25^, cell-extrinsic perturbations ^26^ or sheer chance ^13^ remains unclear. To identify the most highly fit variants we first need to understand how mutation, genetic drift and differences in fitness (selection) combine to produce the spectrum of VAFs observed in CH.

Here, using insights from evolutionary theory, we analyse the VAF spectra of somatic mutations detected in the blood from ∼50,000 individuals to tease apart the effects of mutation, drift and selection. Using single blood sample ‘snapshots’ across many individuals, we quantify the fitness advantages of key pathogenic single-nucleotide variants (SNVs) as well as the spectrum of fitness effects (‘fitness landscape’) of the most commonly mutated driver genes. Using this framework we are able to highlight a number of potentially targetable variants that, while having a low mutation rate and being relatively rare, we estimate to be highly fit and therefore potentially pathogenic. The spectrum of fitness effects in common driver genes is highly skewed: most variants confer either weak or no fitness advantage, but an important minority are fit enough to overwhelm the bone marrow over a human lifespan. We show that positive selection, not drift, is the major force shaping CH and its age dependence ^9,27–29^. Taken together, CH data is consistent with a remarkably simple picture of stem cell dynamics in which HSCs stochastically acquire mutations at a constant rate, which then exponentially expand throughout life.

## Results

### The VAF distribution from ∼50,000 individuals

We analysed VAF measurements for somatic variants in the blood from ∼50,000 blood-cancer-free individuals from nine publicly available blood sequencing datasets ^7–15^ (Supplementary Information 1). VAF measurements in bone marrow and peripheral blood show good concordance ^30^ and so peripheral blood VAF measurements are used as a proxy to reflect clonal composition at the level of the bone marrow HSCs. The nine studies included in our analysis varied in their number of participants and sequencing depth (Figure 1a). Most large-scale studies were limited by standard sequencing error rates and were only able to detect VAFs *>* 3% ^7, 8^ while smaller studies, using error-correcting techniques, were able to detect VAFs as low as 0.03% ^10,12,15^. VAFs varied by over three orders-of-magnitude across individuals, even within the same gene, as exemplified by DNMT3A, the most commonly mutated CH gene (Figure 1b). The density of variants was greatest at low frequencies, with individuals >65 years old almost guaranteed to harbour at least one variant at VAFs*>* 0.03%. Variants were observed far more frequently at certain sites (e.g. R882 hotspot codon, red data) and were almost exclusively putatively functional (nonsynonymous and frameshifts) with synonymous variants being rare and restricted to low VAFs.

**Figure 1.**
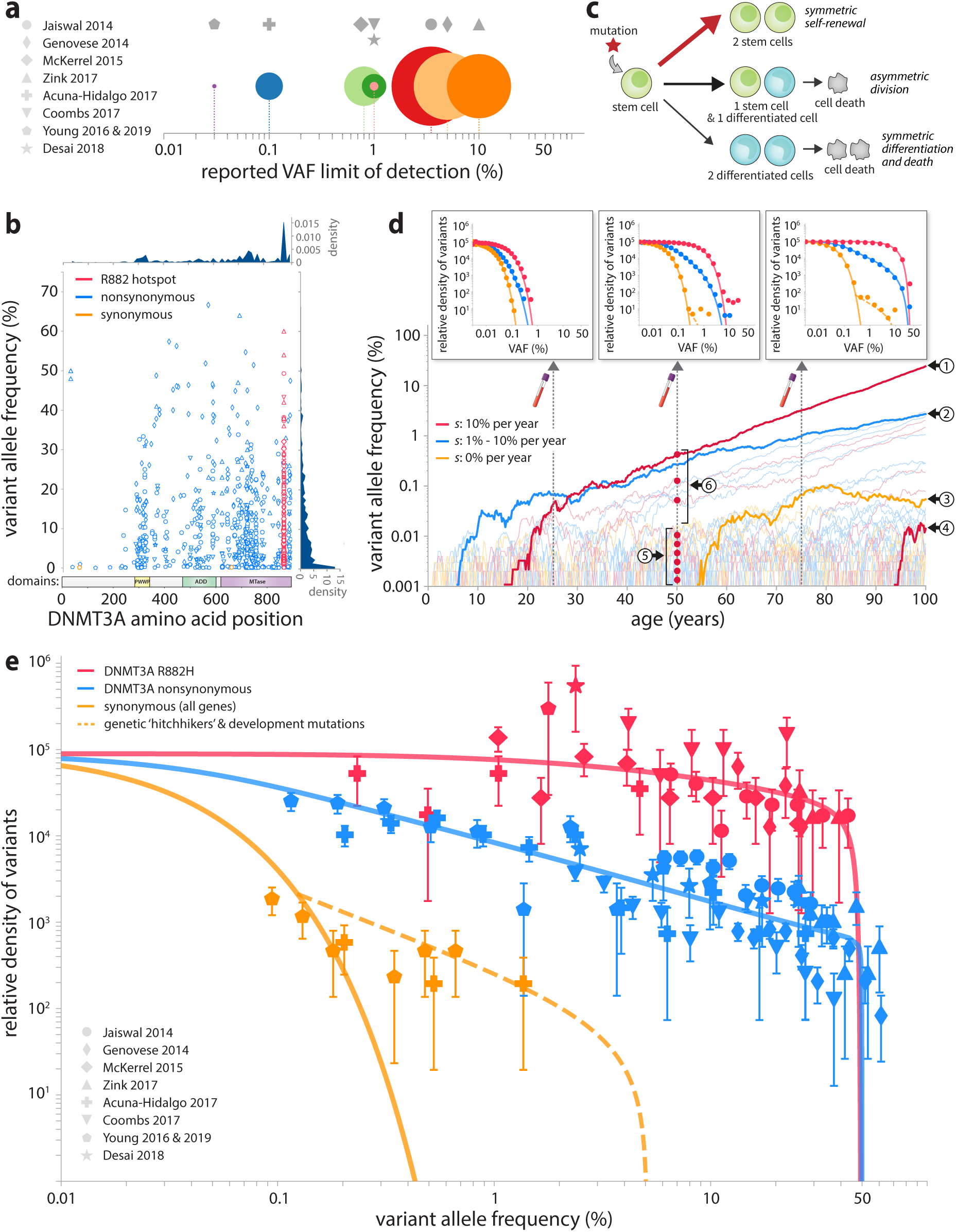
(**a**) Studies used in this analysis varied in number of participants (circle area) and reported VAF detection thresholds. (**b**) The density of variants in DNMT3A varies widely by VAF (> 3 logs) and position in the gene. (**c**) A branching model of HSC dynamics. Mutations with a positive fitness effect (star) cause an imbalance in stochastic cell fates towards symmetric self-renewal (red arrow) resulting in clonal expansions. (**d**) Simulations of HSC populations, using the branching model, show how differences in fitness effect and age produce VAF spectra (insets) in close agreement with observed data (shown in Figure 1e). Numbered features explained in main text. (**e**) Plotting all VAF measurements of DNMT3A variants as log-binned histograms normalised by mutation rates (data points) demonstrates the consistency with the theoretical predictions of the branching model (lines). The theoretical predictions account for a distribution of ages in the studies. The density of high-frequency synonymous variants is consistent with the predicted density of genetic hitchhikers and early developmental mutations (dashed orange line, Supplementary Information 7). Error bars represent sampling noise.

### A branching model of stem cell dynamics

To reveal the relative contributions of genetic drift, mutation rate differences, and cell-intrinsic fitness effects on the observed variation in VAFs, we considered a simple stochastic branching model of HSC dynamics based on classic population genetic models ^31–35^, adapted to include a spectrum of ages and fitness effects (Supplementary information 2). The model is of an HSC population of *N* diploid cells which stochastically self-renew or differentiate symmetrically or asymmetrically (Figure 1c). Mutations are acquired stochastically at a constant rate *µ* per year. The fate of a new mutation depends on its influence on stochastic cell fate decisions via a ‘fitness effect’, *s*, which is the average growth rate per year of that variant relative to unmutated HSCs. Neutral mutations (*s* = 0), do not alter the balance between self-renewal and differentiation and thus either rapidly go extinct or grow slowly, remaining at low VAFs (orange trajectories Figure 1d). Beneficial mutations (*s >* 0) bias cell fates towards self-renewal (red arrow Figure 1c) and, provided they escape stochastic extinction, eventually grow exponentially at rate *s* per year (red and blue trajectories, Figure 1d).

Variants with a high fitness effect or those acquired early in life are expected to reach high VAFs (trajectories 1 & 2, Figure 1d), whereas variants with a low fitness effect or those acquired late in life are restricted to low VAFs (trajectories 3 & 4, Figure 1d). This variation in both the age and fitness effect of variants produces a characteristic spectrum of VAFs measured in a single blood sample (insets, Figure 1d). How these distributions change over a human lifespan is determined by the fitness effect of variants (*s*), their mutation rate (*µ*) and the population size of HSCs (*N*) according to the following expression for the probability density as a function of *l* =log(VAF) (Supplementary information 2):

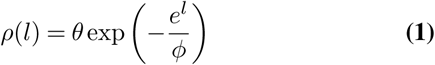

where *l* = log(VAF), *θ* = 2*Nτµ* and 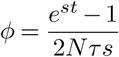

To develop an intuition for the two key features of this distribution, consider variants with a fitness advantage entering the HSC population uniformly at a rate *θ* and growing exponentially. The exponential growth means variant trajectories, plotted on a log-VAF scale, are uniformly spaced straight lines (circle 5, Figure 1d), producing a flat density with y-intercept of *θ*. Dividing the density of variants by the mutation rate (measured per year), the y-intercept therefore provides an estimate for *Nτ* (Figure 1d insets, Supplementary Information 3) where *τ* is the average time between self-renewing HSC divisions (in years). Because the age of the oldest surviving variant cannot exceed the age of the individual, there is a characteristic ‘maximum’ VAF, *ϕ*, a variant can reach, which increases with fitness effect, *s*, and age, *t*. To reach VAFs*> ϕ* requires a variant to both occur early in life and be anomalously lucky, which is unlikely. Therefore, the density falls off exponentially for VAFs*> ϕ* (circle 6, Figure 1d). The sharp density fall-off at 50% VAF is because even a variant that is present in a very large proportion of total HSCs will tend towards 50% VAF due to the cells being diploid.

### Haematopoietic stem cell number and division times

To infer HSC numbers and test the predictions of our model we plotted log-VAF distributions for SNVs from all the studies together. Studies differed in their number of participants as well as their panel ‘footprint’, both of which affect the number of variants detected. Therefore, in order to combine the data from all the studies, we normalised the number of observed variants by their study size and total study-specific mutation rate (for variant or gene of interest), controlling for trinucleotide contexts of mutations (Supplementary Information 4). For a given specific position in the genome, mutation rates are low enough that, over a human lifespan, clones acquiring multiple driver-mutations are rare and thus variants uniquely mark clones (Supplementary Information 5).

We focus first on mutations in the gene DNMT3A (Figure 1e). The most commonly observed variant in DNMT3A is R882H (red data). Because this single variant is expected to confer the same fitness effect, it is a useful first check on the model. Consistent with our predictions, the density of R882H variants is flat over almost the entire frequency range (VAFs<15%) with a y-intercept of *Nτ ≈τ* 100, 000 ± 30, 000 years (Supplementary Figures 8 and 10). Encouragingly, this number is in very close agreement with that inferred from single HSC phylogenies ^36^. An important point to note is that ours and other population genetic analyses can only infer the combination *Nτ* and not *N* or *τ* separately, but, combined with estimates of HSC symmetric division rates of 0.6– 6 per year ^36–38^, this data suggests there are between 60,000– 600,000 HSCs maintaining the peripheral blood.

To validate our estimates for *Nτ*, we turned to the distribution of all synonymous variants (orange data, Figure 1e). Because synonymous variants are generally expected to be functionally neutral, the characteristic VAF of the biggest synonymous variants (*ϕ*) increases only linearly with age, as it is driven by drift alone (see eqn.1). This provides a crucial validation of the model since it predicts that the majority of synonymous variants should be found at very low VAFs. Quantitatively, if our inferred value of *Nτ ≈* 100, 000 years from R882H variants is correct, it would predict that the majority of synonymous mutations should be restricted to VAFs below *ϕ* = *t/*2*Nτ ≈* 0.025% at age 50. This prediction broadly agrees with the data, where the maximum likelihood inferred *ϕ* ≈ 0.03 ± 0.01% (Supplementary Information 6). This internal consistency check indicates that both synonymous and DNMT3A R882H variants point toward similar values of *Nτ*. Synonymous variants with VAFs ≫ *ϕ* are rare (orange dashed line, Figure 1e) and are consistent with having hitchhiked to high frequencies on the back of an expanding clone that had already acquired a fit variant (Supplementary Information 7), although it is also possible a handful are developmental in origin, or have a functional consequence themselves e.g. due to codon usage bias, or are in fact non-synonymous in an alternatively spliced transcript.

### The fitness landscape of clonal haematopoiesis

Because the characteristic maximum VAF, *ϕ*, depends on the fitness effect, *s*, by estimating *ϕ* from the VAF spectrum, we can infer a variant’s fitness. We illustrate this approach using DNMT3A R882H variants. As predicted by the model, the density of R882H variants does indeed begin to fall off exponentially for VAFs>12% (red data, Figure 1e and Supplementary Information 6). This suggests that R882H variants provide HSCs with a large selective advantage (*s ≈* 15 ± 1% per year) since, over the course of ≈ 55 years (mean age across all studies), they have expanded to VAFs ≈ 12%, although some have reached VAFs as high as 50%.

To reveal the fitness landscape of other highly fit and possibly pathogenic variants we applied this analysis to each of the 20 most commonly observed variants across all studies (Figure 2a). Variants in the spliceosome genes SF3B1 and SRSF2 are some of the fittest in CH, with fitness effects as high as *s* ≈ 23% per year, but are relatively rare due to low mutation rates. DNMT3A R882H is the most common CH variant, not because it is the most fit, but because it is both highly fit and has a high mutation rate due to its CpG context. The DNMT3A R882C variant is in fact significantly fitter than R882H (*s ≈* 19 ± 1% vs ≈ 15 ± 1% per year) and is only observed less frequently because of its lower mutation rate (Supplementary Information 6). The potential of our analyses is underscored by the GNB1 K57E variant. While this variant has received little attention in CH, it is in fact highly fit and, importantly, strongly associated with myeloid cancers and potentially targetable ^39^.

**Figure 2.**
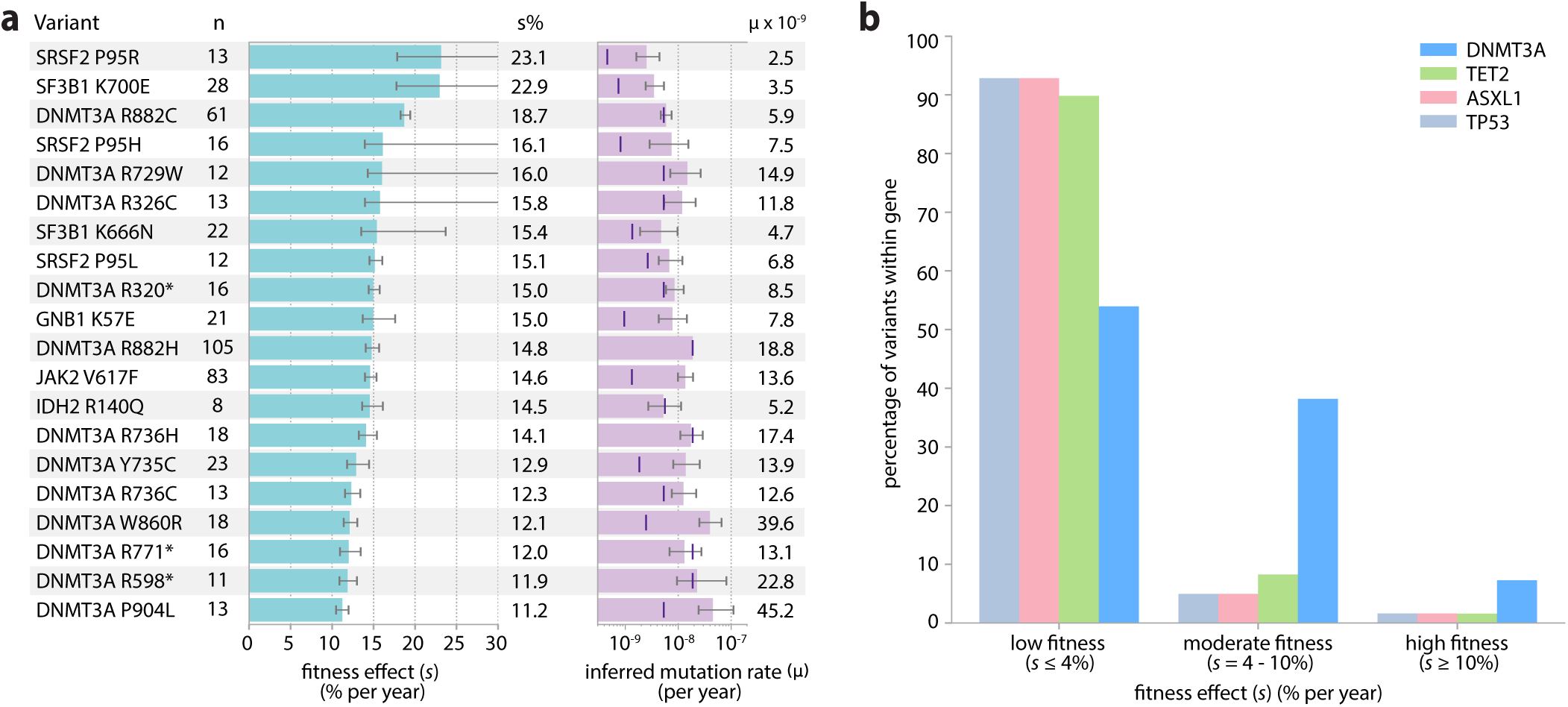
The fitness landscape of clonal haematopoiesis variants and genes. (**a**) Inferred fitness effects and mutation rates for the top 20 most commonly observed CH variants. Error bars represent 95% confidence intervals. Purple vertical lines indicate site-specific mutation rates inferred from trinucleotide context (Supplementary Information 4). (**b**) The distribution of fitness effects of nonsynonymous variants in key CH driver genes, inferred by fitting a stretched exponential distribution and dividing this up into three fitness classes (low, moderate and high) (Supplementary Information 6). These distributions reveal many low fitness and few high fitness variants (Table inset). Over a human lifespan, variants with fitness effects <4% expand only a modest factor more than a neutral variant (‘low fitness’), variants with 4%-10% per year expand by substantial factors (‘moderate fitness’) and variants with fitness effects*>* 10% per year can expand enough to overwhelm the marrow (‘high fitness’).

To reveal the overall fitness landscapes of key CH driver genes we considered the VAF distribution of all nonsynonymous variants in each of the genes DNMT3A, TET2, ASXL1 and TP53 (Figure 2b). For DNMT3A, the density of nonsynonymous variants at low VAFs is broadly consistent with the same *Nτ ≈* 100, 000 years inferred from R882H variants (Figure 1e, blue data). However, with increasing VAF the density of variants declines, consistent with a spectrum of *ϕ* and thus a spectrum of fitness effects. Performing a maximum likelihood fit to a family of stretched exponential distributions, we found that the spectrum of *Φ* fitness effects for nonsynonymous variants in DNMT3A is very broad with ≈ 46% of variants conferring moderate to high fitness effects (*s >* 4% per year, Figure 2b, Supplementary Information 6). In contrast, the genes TET2, ASXL1 and TP53 have a spectrum that is more skewed towards low fitness effects with only ≈ 7–10% of all possible nonsynonymous variants in these genes conferring moderate or high fitness effects. These distributions highlight that, in these CH genes, most nonsynonymous variants have a low enough fitness that they are effectively neutral, while an important minority expand fast enough to overwhelm the marrow over a human lifespan.

### Age-dependence of clonal haematopoiesis

A key prediction of the model is that, because variants enter the HSC population at a constant rate, the apparent prevalence of a specific variant, at a defined sequencing sensitivity, is predicted to increase roughly linearly with age, at rate 2*Nτ µs* (Supplementary Information 8). We confirmed this prediction using DNMT3A R882H and R882C variants which, when combined, had enough data to be broken-down by agegroup (Supplementary Figure 15). In agreement with predictions, the age-prevalence of these variants does increase linearly with age, consistent with the age-dependence of CH being driven by the expansion of clones which become more detectable at older ages. The rate of this increase provides an independent way to validate estimates of fitness effects and, in this case, the rate of increase is consistent with a fitness effect of *s ≈* 14% per year, which is in good agreement with estimates inferred from the VAF distribution (Figure 2a).

By inferring the spectrum of fitness effects across ten of the most commonly mutated CH genes, we can predict how common CH will be as a function of both age and sequencing sensitivity (Figure 3a and Supplementary Information 9). With sensitive enough sequencing (VAFs; ≳ 0.01%), CH variants will be common even in young adults and almost ubiquitous in people aged over 50, consistent with recent ultra-sensitive sequencing studies ^10, 15^. Our framework also enables us to predict the emergence of clones harbouring multiple driver mutations. While this depends on the cooperativity between mutations, under the assumption of additive fitness effects we predict that *<* 10% of individuals aged 80 years old will harbour clones with multiple-driver mutations (Supplementary Information 5).

**Figure 3.**
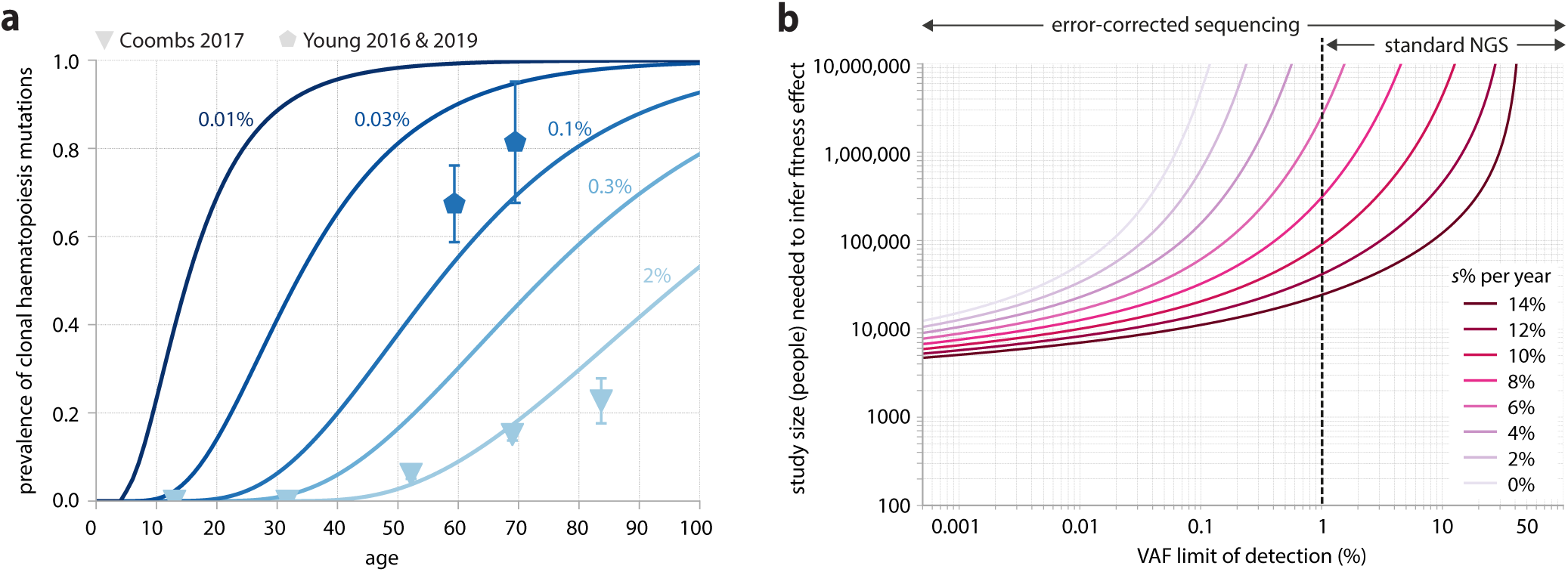
(a) Predicted prevalence of CH mutations as a function of age for different detection thresholds. Prevalence is predicted for individuals to have acquired at least one variant within 10 of the most commonly mutated CH genes (DNMT3A, TET2, ASXL1, JAK2, TP53, CBL, SF3B1, SRSF2, IDH2 and KRAS), taking in to account the distribution of fitness affects across these genes (Supplementary Note 9). The actual prevalence of variants within these genes, as a function of age, is shown for Young 2016 & 2019 (hexagons, VAF limit of detection 0.1%) and Coombs 2017 (triangles, VAF limit of detection 2%). **(b)** Study size required to accurately quantify different fitness effects (coloured lines) for individual variants, with an average site-specific mutation rate of 1.3 *×* 10^−9^as a function of sequencing sensitivity (VAF limit of detection).

## Discussion

### A simple framework explains clonal haematopoiesis

Analysing the VAF spectra from nine publicly available clonal haematopoiesis datasets in light of evolutionary theory points to a remarkably simple and consistent picture of how HSC population dynamics shape the genetic diversity of blood. The very wide variation in VAFs observed between people is largely caused by the combined effects of chance (when a mutation arises) and fitness differences (how fast they expand). While cell-extrinsic effects are likely crucial in specific contexts such as chemotherapy ^11,26,40,41^ and acute infection ^42, 43^, the data from healthy individuals are quantitatively consistent with cell-intrinsic differences in fitness effects being the major determinants of the variation.

While it might seem surprising that a simple model captures many quantitative aspects of CH data, more complex scenarios yield the same effective model for the multi-year development of CH (although *N* and *τ* have more complex meanings). These include models with HSCs switching between active and quiescent states, and models with progenitors occasionally reverting to HSCs. But, there are important observations that the model cannot fully explain, including a considerably broader than expected distribution in the number of variants observed in different individuals, although this could be attributed to variations in mutation rates across individuals or environment-specific effects. Distinguishing between these scenarios will likely require longitudinal data and is an important area for future work.

### In haematopoietic stem cells, fitness dominates drift

The relative roles of mutation, drift and selection in shaping the somatic mutational diversity observed in human tissues has been the subject of much recent debate, especially regarding the conflicting interpretations from ‘dN/dS’ measures ^1,5,44^ and clone size statistics ^34,45,46^. In blood, the two measures are in quantitative agreement; non-synonymous variants are under strong positive selection and most synonymous variants fluctuate via neutral drift.

Our inference of the large HSC population size (*Nτ ≈* 100,000 years) has an important interpretation: on average it would take an absurd 100, 000 years for a variant to reach VAFs of 50% by drift alone and >2000 years to be detectable by standard sequencing (1%). Therefore the vast majority of CH variants reaching VAFs*>* 0.1% over a human lifespan likely do so because of positive selection. However, this is not to say that variants with VAFs*<* 0.1% are not potentially pathogenic. Indeed, most highly fit variants exist at low VAFs simply because not enough time has yet passed for them to expand, although they are less likely to acquire subsequent driver mutations whilst they are at low VAFs.

### Over 3000 variants confer moderate to high fitness

By considering the VAF spectrum across ten of the most commonly mutated CH genes, we have inferred that mutations conferring fitness effects *s >* 4% per year occur at a rate of ≈ 4.5 *×*10^−6^ per year (Supplementary Information 9). Given the average site-specific mutation rate in HSCs is 1.3 *×* 10^−9^ per year (Supplementary Information 4), this implies there are; ≳ 3000 variants within these genes conferring moderate to high selective advantages. While there is direct evidence from longitudinal data ^18^ and indirect evidence from age-prevalence patterns (Supplementary Information 8) that variants at many of these sites expand at a roughly constant rate, others, notably JAK2 V617F, might exhibit more complex dynamics given the small exponential growth rates observed in longitudinal data ^47^.

The variants commonly observed in CH are not necessarily the most fit, but are both sufficiently fit and sufficiently frequently mutated. To reveal variants that are infrequently mutated, yet potentially highly fit, we considered all variants in DNMT3A, TET2, ASXL1 and TP53 that were detected at least twice across all nine studies and estimated their fitness effects by determining what fitness effect would be needed to produce the number of observed variants (Supplementary Information 10). While the lack of data at infrequently mutated sites and the crudeness of this counting method necessarily leads to large uncertainties, there appears to be at least some highly fit yet infrequently mutated variants which, while individually rare, could be collectively common (Supplementary Information 10).

How large would a study need to be to exhaustively sample all moderate and highly fit variants and quantify their fitness effects? Given the average site-specific mutation rate of 1.3 *×* 10^−9^ per year (Supplementary Table 4), a comprehensive map between mutation and fitness effect for all sites that confer a selective advantage large enough to expand significantly over a human lifespan (*s >* 4%) could be achieved with the current sample size by increasing sequencing sensitivity to detect variants at VAFs*>* 0.03% (Figure 3b). However, because sites can mutate at rates as low as *µ* ∼ 10^−10^ / year (Supplementary Table 4) to quantify all variants, even rare ones, would require both a 6-fold increase in sample size, as well as sequencing sensitivities as low as 0.01% VAF (Supplementary Information 11). Nonetheless, even with small study sizes, there are major advantages to being sensitive to very low VAFs ^10,12,15^, particularly in relation to synonymous variants, which, when grouped together, provide important information on *Nτ* and genetic hitchhikers (Figure 1e).

The absence of variants in known AML drivers such as FLT3 and NPM1, across the nine studies, suggests that mutations in these genes do not confer an unconditional selective advantage to HSCs, consistent with studies in mice and humans showing that they are late-occurring and possibly cooperating mutations necessary for transformation to AML ^20, 23^.

### Future directions

CH has associated risks with cardiovascular disease ^7, 48^ and progression to blood cancers ^7,8,14,18^, and has important consequences in the study of ctDNA ^49, 50^, aplastic anaemia ^51^, response to chemotherapies ^52, 53^ and bone marrow transplant ^40,54,55^. A major challenge is to develop a predictive understanding of how variants and their VAFs affect disease risk. Pioneering recent studies have made great strides in this direction, showing that both gene identify and VAF are predictive of progression to AML ^14, 18^. The framework presented here provides a rational basis for quantifying the fitness effects of these variants and understanding VAF variations. Combining this framework with studies that longitudinally track individuals over time will shed light on how these initiating mutations acquire further mutations that drive overt disease. More sensitive sequencing techniques, broader sampling of the genome (e.g. regulatory regions) and the study of environmental factors that alter the fitness of mutations, will improve our quantitative understanding of native human haematopoiesis and accelerate the development of risk predictors.

## ACKNOWLEDGEMENTS

We thank all members of the Blundell, Fisher and Druley labs. We thank Sasha Levy, Ivana Cvijovic, Dmitri Petrov, Ben Simons, Moritz Gerstung, Brian Huntly, Inigo Martincorena, Ross Levine, Sidd Jaiswal and Ravi Majeti for helpful comments. C.J.W and J.R.B are funded by the CRUK Cambridge Centre and CRUK Early Detection Programme. A.P. is supported by the National Science Foundation GRFP. D.S.F and J.R.B. are supported by the Stand Up to Cancer Foundation and the National Science Foundation via PHY-1545840.

## Supplementary Note 1: Data used in analysis

### Studies and samples included in our analysis

A total of nine publicly available blood sequencing datasets were included in our analysis. These studies were those whose panel ‘footprint’ we felt could be reliably determined from their published information and were either large studies (≥1000 participants) or used deep-sequencing methods (VAF detection limit ≤1%) (Table 1). The size of the panel ‘footprint’ (Supplementary Information 4), affects the number of variants detectable and so having this information was essential to enable meaningful comparisons of VAF densities across studies.

**Table 1.** Details of studies included in our analysis. ECS: Error-corrected sequencing, HPFS: Health Professionals Follow-Up Study, MSKCC: Memorial Sloan Kettering Cancer Center, NBS: Nijmegen Biomedical Study, NHS: Nurses Health Study, smMIPs: Single Molecule Molecular Inversion Probes, T2DM: Type 2 Diabetes Mellitus, UKHLS: UK Household Longitudinal Study, WES: Whole Exome Sequencing, WGS: Whole Genome Sequencing, WHI: Women’s Health Initiative, WTCCC: Wellcome Trust Case Control Consortium

| Study | Study size | Participants | Haematological characteristics | Age range | Sex | Sequencing method | Reported limit of SNV detection | Participants included in our data analysis |
| --- | --- | --- | --- | --- | --- | --- | --- | --- |
| **Jaiswal 2014<sup>7</sup>** | 17182 | Population-based cohorts (15801 cases/controls from T2DM association studies, 1381 from Jackson Heart study) | Unselected for haematologic phenotype. | 19-108 | 51% female, 49% male | WES | 3.5% | All (17182) |
| **Genovese 2014<sup>8</sup>** | 12380 | Swedish individuals (6245 controls, 4970 schizophrenia, 1165 bipolar disorder) | Unselected for haematologic phenotype. | 19-93 | 49% female, 51% male | WES | 5% | All (12380) |
| **McKerrel 2015<sup>9</sup>** | 4219 | 3067 UK blood donors (WTCCC), 1152 UKHLS participants | Unselected for haematologic phenotype. | 17-98 | ? | Barcoded multiplex PCR of mutational hotspots. | 0.8% | All (4219) |
| **Young 2016<sup>10</sup>** | 20 | Population-based cohort (NHS) - all had blood sample at 2 time-points | Unselected for haematologic phenotype, but no history of cancer or major chronic disease. | 51-74 | 100% female | ECS (adapted Illumina TruSight Myeloid Panel). | 0.03% | All (only 1 time-point) |
| **Zink 2017<sup>13</sup>** | 11262 | Icelanders participating in various disease projects at deCODE genetics. | Excluded from study if diagnosis of haematological malignancy before or within 6 months after blood sample. | <10 - >100 | 55% female, 45% male | WGS and then targeted resequencing on Illumina TruSight Myeloid Panel of 72 individuals with CH. | 10% (WGS) | All (11262) (only WGS) |
| **Acuna-Hidalgo 2017<sup>12</sup>** | 2006 | Population-based cohort (NBS). | Unselected for haematologic phenotype. | 20-69 | 50% female, 50% male | smMIPs | 0.1% | All (2006) |
| **Coombs 2017<sup>11</sup>** | 5649 | Patients with non-haematologic cancers at MSKCC. | Excluded from study if active haematologic cancer or precursor conditions. | <1-98 | 51% female, 49% male | Hybridization capture-based sequencing assay encompassing all protein-coding exons of 341 or 410 cancer-associated genes (MSK-IMPACT). | 1% | Only those who were chemotherapy-naïve and radiotherapy-naïve (1591) |
| **Desai 2018<sup>14</sup>** | 424 | Individuals who developed AML and controls (~1:1 ratio) within WHI cohort - >200 with samples from ≥2 time-point. | Controls excluded if history of haematologic disorder or AML during study. | 51-79 | 100% female | Targeted sequencing using custom capture probes (Nimblegen). | 1% | Only controls (181) - only 1 time-point (base-line sample) |
| **Young 2019<sup>15</sup>** | 103 | Individuals who developed AML and controls (2:1 ratio) within NHS and HPFS cohorts - all NHS participants had sample from 2 time-points. | Controls excluded if history of cancer (other than non-melanoma skin cancer). | 48-79 | 45% female, 55% male | ECS (adapted Illumina TruSight Myeloid Panel) | 0.03% | Only controls (69) (only 1 time-point included) |
| **TOTAL INCLUDED IN OUR ANALYSIS:** |  |  |  |  |  |  |  | 48910 |

All of the participants in Coombs 2017^11^ were individuals who had been diagnosed with a non-haematological malignancy. We only included in our analysis the individuals that were both chemotherapy-naive and radiotherapy-naive. Two of the studies, Desai 2018^14^ and Young 2019^15^, were nested case-control studies designed to assess the relationship between clonal haematopoiesis and risk of progression to AML. We only included the control individuals in our data analysis. Two of the studies, Young 2016^10^ and Young 2019^15^, reported replicate measurements for their VAFs. We required a variant to be detected in both replicate samples to be called and the average of the replicate values was taken as the VAF at that time-point. Three of the studies, Young 2016^10^, Young 2019^15^ and Desai 2018^14^, reported variants in participants from more than one time-point. We only included variants detected in the first blood sample for these studies.

### Data trimming below study-specific limits of detection

Only single nucleotide variants (SNVs) were included in our analysis, due to mutation rate uncertainties for other classes of mutation. While studies generally reported their VAF detection threshold, this is typically determined by a predetermined false positive rate, at which false negative rates could be substantial. To estimate where false negative rates were beginning to have a substantial effect on the data, we used variants in DNMT3A (which had the most data) and chose a threshold VAF below which the density began to decline (Supplementary Figure 1, Table 2). Variants were not included in our analysis if their VAF was below this study-specific VAF threshold. The same threshold was also used for trimming all other variants reported by that study.

**Supplementary Figure 1.**
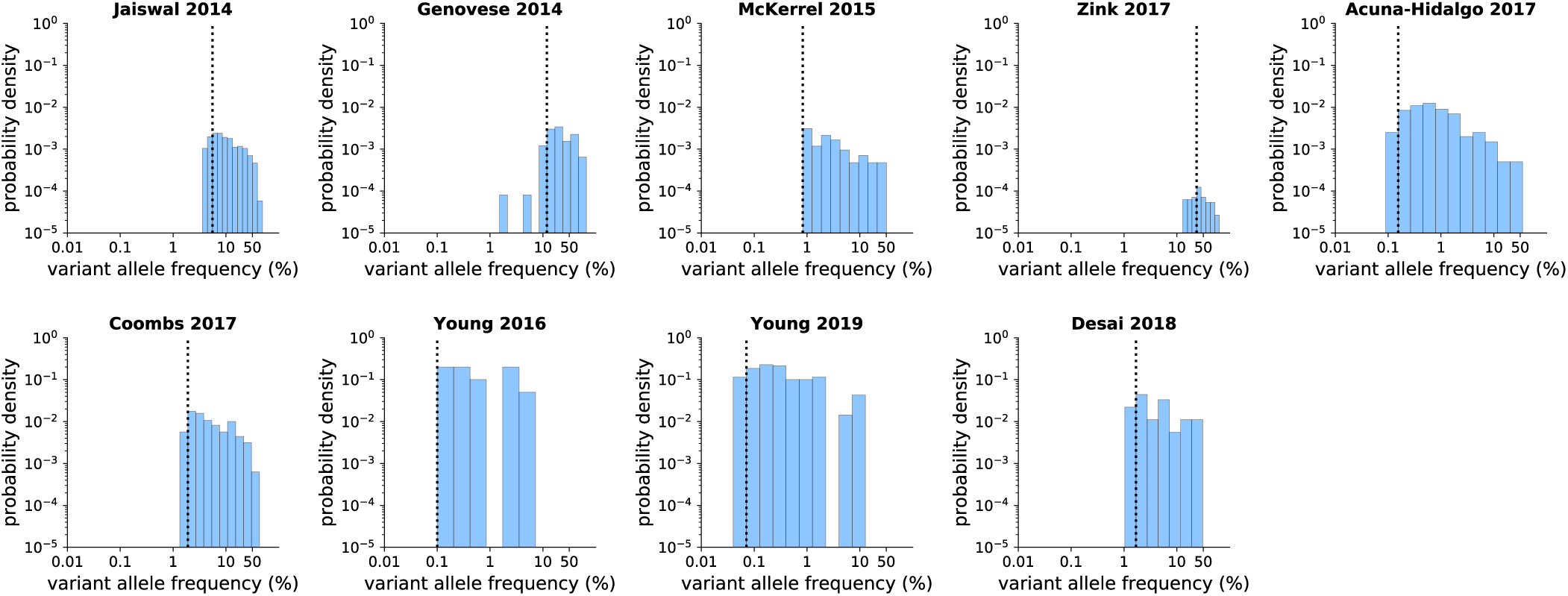
Data trimming of DNMT3A variants. Vertical dashed lines on the probability density histograms indicate the VAF below which the density of DNMT3A variants starts to fall off and thus likely represents the study’s limit of reliable variant detection. DNMT3A variants at VAFs lower than this cut-off were not included in our data analysis and this cut-off was also used for trimming all other variants reported by that study. The height of the histograms are representative of the sequencing depth.

**Table 2.** Study-specific limits of detection used for data trimming. The VAF limit of SNV detection reported by each study is shown in comparison to the study-specific threshold below which we trimmed each study’s data.

| Study | Reported VAF limit of SNV detection | VAF threshold used for data trimming |
| --- | --- | --- |
| Jaiswal 2014<sup>7</sup> | 3.5% | 5.56% |
| Genovese 2014<sup>8</sup> | 5% | 11.94% |
| McKerrel 2015<sup>9</sup> | 0.8% | 0.8% |
| Zink 2017<sup>13</sup> | 10% | 23.63% |
| Coombs 2017<sup>11</sup> | 1.00% | 1.90% |
| Acuna-Hidalgo 2017<sup>12</sup> | 0.1% | 0.15% |
| Young 2016<sup>10</sup> | 0.03% | 0.10% |
| Desai 2018<sup>14</sup> | 1% | 1.69% |
| Young 2019<sup>15</sup> | 0.03% | 0.07% |

## Supplementary Note 2: Theory for the variant allele frequency distribution

We consider a continuous time branching process for haematopoietic stem cells (HSCs). In the absence of mutation, in an interval *dt* a single HSC can (i) divide symmetrically producing two stem cells with probability *λr*(1 + *s/*2)*dt*, (ii) divide symmetrically producing two terminally differentiated cells with probability *λr*(1 – *s/*2)*dt*, (iii) divide asymmetrically producing one HSC and one terminally differentiated cell with probability *λ*(1 – *r*)*dt*. Where *λ* is the HSC division rate, *r* is the fraction of cell divisions that are symmetric and *s* is a bias towards self-renewal and is equivalent to a selective advantage. We are interested in the probability distribution of clone sizes *P*_*n*_ which can be calculated from a master equation using the transition probabilities *T* (*n –* 1 → *n*) = (*n –* 1)*λr*(1 + *s/*2)*dt, T* (*n* + 1 → *n*) = (*n* + 1)*λr*(1 – *s/*2)*dt*, and *T* (*n* → *n*) = 1 – 2*nλrdt*. Taking the continuous *n* limit and rescaling units of time to be measured in units of symmetric cell divisions *τ* = *λrt* gives the Fokker-Planck equation for the dynamics of *ρ*(*n, t*)

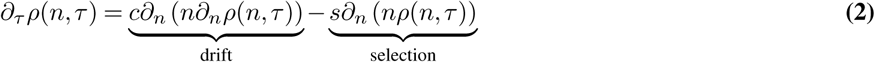

with the initial condition *ρ*(*n*, 0) = δ(*n – n*_0_) and where *c* = 1. This can be solved using generating functions i.e. taking a Laplace transform by multiplying both sides by exp (–*nx*(*τ*)) and integrating over *n*. This generates a differential equation for *x*(*τ*)

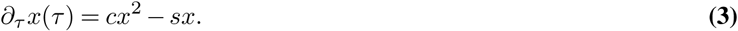

This logistic equation can be solved subject to the initial condition that at *τ* = 0, *M* (*ϕ*) = *e*^*-n*^0^*x*(*τ* =0)^, so using the differential equation to relate *x*(*τ* = 0) to *x*(*τ* = *T*). The solution is

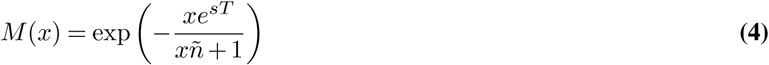

where ñ = (*c/s*) *e*^*sT*^ *–* 1. This can then be inverted either via an inverse Laplace Transform (using steepest descents) or by small expanding *e*^*∈*^ *≈* 1 + *∈* from which one recovers:

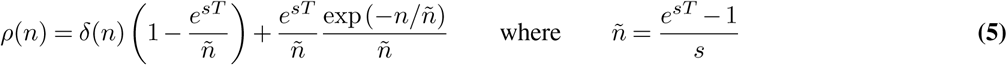

This is a good approximation to the distribution of clone sizes at time *T* given a single-cell was present at initially.

If mutations at a particular position occur at a constant rate *θ* = 2*NU* (the factor of 2 from the fact that there are two possible alleles to mutate), then the clone-size distribution is the convolution of the single mutant distribution (above) with the distribution of times at which they enter (which is uniform). Convolutions are products in the moment generating function hence:

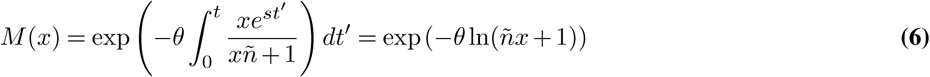

where, since the units of time are measured in symmetric cell divisions, the rate of mutation *µ* must be measured in these same units i.e. the mutation rate at that position per symmetric cell division, which if *r* ≪ 1 could be significantly larger than the mutation rate per cell division. The inverse Laplace Transform can be performed exactly in this case via the residue theorem and yields *ρ*(*n*) = *e*^*-n/ ñ*^*/*(G(*θ*)*n*^1*-θ*^). For a given position the value of *θ* ≪ 1 and thus the expression for the density of clones with size *n* can be well approximated by

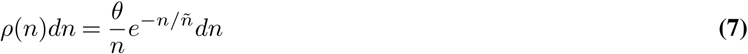

Changing variables so that frequencies are measured on a log-scale *l* = ln(*n/*2*N*) and defining *ϕ* = ñ */*(2*N*) this distribution becomes

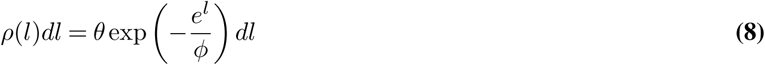

which is the result quoted in the main text. Since the expanding clone itself contributes to the total number of cells, a slightly more accurate expression for the fraction of variant reads is obtained by the change of variables 2*f* = *n/*(*n* + *N*), giving

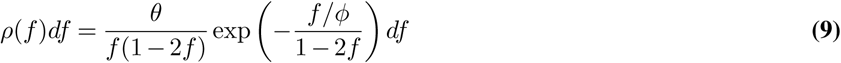

where *ϕ* = ñ */*(2*N*). This result is used in Figures of the main text and for the parameter estimates for *N* and *s* and *ρ*(*s*).

We validated this theoretical prediction by simulating the dynamics of mutations in the gene DNMT3A across 100,000 individuals each with *N* = 10^5^ HSCs and plotted the distribution of VAFs at three different ages (Figure 1d, Supplementary information 3). Because of the 1*/f* divergence at low frequencies and the exponential “cutt-off” at high frequencies, an informative way of visualising the predicted density is to plot it as a function of the log VAF as shown in (Figure 1). Plotting the clone density in this way is desirable because the two key parameters *θ* and *ϕ* separate (Supplementary Figure 2). The estimate for *θ* is simply the y-intercept while *ϕ* is the frequency scale at which the density begins to fall off away from *θ*, and begins to decay exponentially.

### Units

The value of *θ* = 2*NU* in eqn. 9 is the total rate at which new single-mutants are generated per symmetric cell division, where *N* is the number of stem cells and *U* is the haploid mutation rate. However, the rate of symmetric cell division is challenging to measure. The measurable quantity is the mutation rate per year *µ* = *U/τ* where 1*/τ* = *λr* is the number of symmetric divisions per year, hence

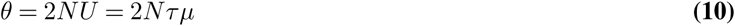

Because *θ* can be estimated directly from the histogram (see Supplementary Information 6) and *µ* can be robustly estimated (Supplementary Information 4), we can directly infer the ratio *Nτ* from the data without making any strong assumptions. This is the number of HSCs multiplied by the HSC symmetric division time (in years). Hence if *Nτ* = 100, 000, this would be 100,000 HSCs dividing once per year or 400,000 HSCs dividing once per 3 months.

The value of *ϕ* = (exp(*st*) – 1)*/*2*Ns* (using time in units of symmetric cell divisions) is the characteristic frequency of extant clones (it is the average size of clones that enter at *t* = 0). If using units of time in years, *a*, we get *ϕ* = (exp(*Sa –* 1)*/*2(*Nτ*)*S* where *S* is the fitness advantage of the clone per year. For neutral mutations, *ϕ* = *t/*2*N* = *a/*2*N τ*, where *a* is the age of the person in years. Since the synonymous *ϕ* and the age *a* are both directly measure-able from the data, the ratio *ϕ/a* provides an independent check on *Nτ*. This means that an internal consistency check is:

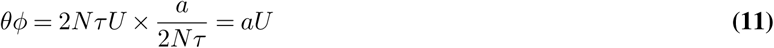

**this provides a way of checking internal consistency without having to make assumptions about** *τ* **or** *N* **independently**.

**Supplementary Figure 2.**
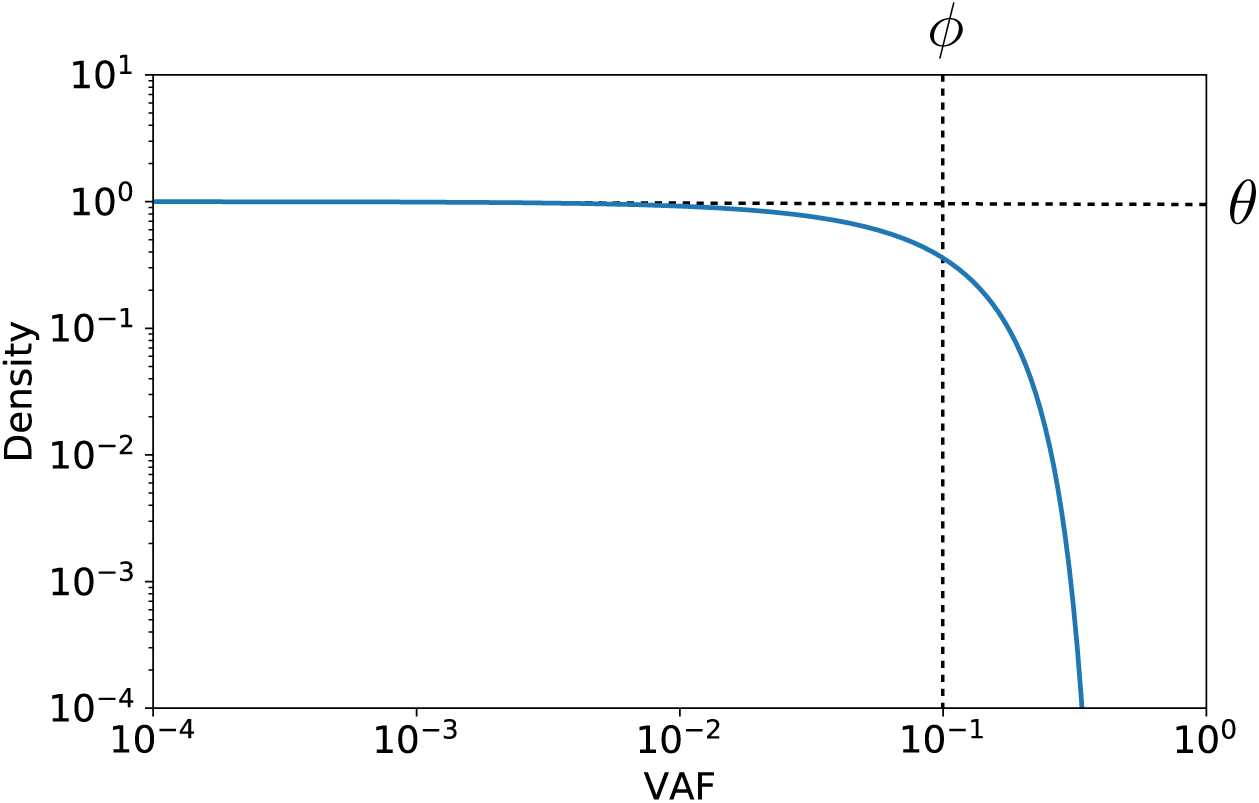
Density of clones as a function of VAF plotted on log-log axes. The universal scaling at low frequencies manifests as a constant density below *ϕ*, while above *ϕ* the decline is exponential.

## Supplementary Note 3: Stochastic Simulations of Clone Dynamics

We simulated the evolutionary dynamics using a custom written stochastic process implemented in Python. Briefly:

- The code simulates a population (“ `population` ”), which is a list of all unique clones, and their abundances, present in the population at a given time.
- clones are tuples composed of “(`clone_ID`, `clone_size`)” where `clone_size` is the integer number of cells comprising the clone, and, where `clone_ID` is a list of (`mutation_ID`, `fitness_effect`) pairs for all unique mutations that have accumulated in that clone.
- `mutation_ID` uniquely labels each independent mutation to have entered the population and is updated via a counter called `last_mutation_ID`, thus a `mutation_ID` = 5 means that was the 5th mutation to occur in the population.
- `fitness_effect` is the selective effect, *s*, of the mutation and is randomly drawn from the distribution of fitness effects `DFE`, which can be varied (see below). Fitness effects combine additively i.e. a clone with two mutations with fitness effects *s*_1_, *s*_2_ would have a fitness effect *s*_1_ + *s*_2_
- The dynamics are implemented using two functions `mutate` (which generate new clones) and `select` (which modifies the clone sizes of existing clones) in discrete time steps (`dt=0.1`) where units of time are measured in years.
- The function `mutate` creates new clones by querying each clone in the current population and determining the number of daughter clones each gives rise to by drawing a Poisson random variate with mean `clone_size × u × dt`, where `u` = 3 × 10^−6^ is the mutation rate for each cell in the clone. New clones are added to the list of all clones with an initial `clone_size` = 1.
- Clones fitness effects are drawn from a `DFE` (distribution of fitness effects) which we typically set to *P* (*s* = 0) = 1*/*3 and *P* (*s >* 0) = 2*/*3 though the precise distribution of the *s >* 0 depends on context. For a single site the *s >* 0 mutations are a single value. For a gene (e.g. DNMT3A) these can be drawn from a distribution e.g. an exponential *ρ*(*s*) ∝ exp (–*s/d*) over a range 0 *< s < s*_*max*_.
- The function `select` updates the clone sizes of existing clones determining the difference in the number cell-births and cell-deaths occurring in that clone. In the time interval `dt` these are calculated via

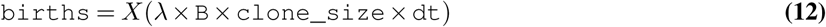

where *X*(*g*) is a Poisson random variate with mean *g* and where *λ* = 5 is the total cell division rate (i.e. we assume 5 HSC divisions per year). Similarly the number of deaths is

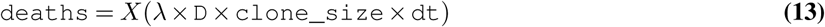 The fitness differences are implemented by allowing birth-rates to depend on the fitness of the mutations in the clone:

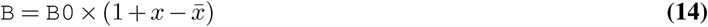

where `D` = `B0` = 0.2. Thus, when there is no fitness advantage 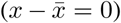 the clone performs neutral drift due to matched birth and death rates i.e. one in every 5 cell divisions results in a self-renewal, and another 1 in 5 results in a symmetric differentiation event. The fitness advantage (which can be negative) is determined by calculating the fitness of the clone, *x*, as the sum of the fitness effects of all mutations it has acquired (*x* = ∑_*i*_ *s*_*i*_) and determining its advantage over the mean fitness, 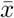 of all clones in the population using 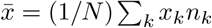 where *n*_*k*_ is the clone size and *k* indexes all clones in the population. The mean fitness ensures the total population of HSCs remains constant (up to small stochastic fluctuations). In the simulations outlined in the main text, we do not implement a mean fitness, meaning that the population of HSCs has the potential to grow. We do this because, for the parameters used, clones rarely grow to take over an appreciable fraction of the population hence there is little difference between using mean-fitness and not.
- The `clone_size` and `population_size` are then recorded at any time required and are used to calculate the variant allele frequency, `vaf`, via `vaf` = `clone_size` */*(2 *×* `population_size`), where the factor of two comes from the fact cells are expected to be diploid.

## Supplementary Note 4: Mutation-rate estimates

Two major studies have published large-scale data on mutations found in single-cell derived HPSCs ^36, 56^. Though we use the data from Lee-Six et al’s study ^36^, there is close agreement between these two studies. Lee-Six et al. performed whole-genome sequencing (to a depth of 15X) on 140 single-cell derived HSC colonies from a healthy 59-year-old male ^36^. They identified 129,582 genome-wide somatic mutations across the 140 colonies which, over 59 years of life, equates to ≈ 15.7 mutations per year per genome (≈ 5.2 *×* 10^−9^ per bp per year). They categorised the observed substitutions into 96 trinucleotide-context specific categories according to the pyrimidine base change and its neighbouring 5’ and 3’ bases (e.g. A[C>A]A). To obtain site-specific mutation rate estimates (per year), for these 96 site contexts, as well as their complementary site contexts (i.e. total 192 site contexts), we normalised by the trinucleotide frequencies (of both sites) across the mappable genome (5.87 10^9^ bp per cell). Trinucleotide frequencies were calculated in R/Bioconductor using the *BioStrings* package ^57^ using *BS.genome.Hsapiens.UCSC.hg19* (Table 3). The normalised number of substitutions was then divided by the number of colonies (140) and by the age of the individual (59), in order to obtain a haploid trinucleotide-context-site-specific mutation rate in units of years (Table 4):

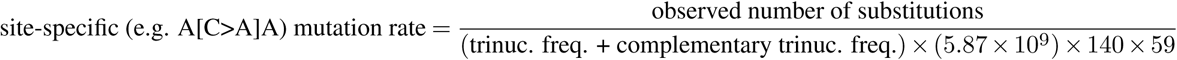

**Table 3.** Trinculeotide frequencies across the genome.

| Site | % of genome | Site | % of genome | Site | % of genome | Site | % of genome |
| --- | --- | --- | --- | --- | --- | --- | --- |
| AAA | 3.8356 | CAA | 1.8905 | GAA | 1.9703 | TAA | 2.0782 |
| AAC | 1.4548 | CAC | 1.5012 | GAC | 0.9439 | TAC | 1.1341 |
| AAG | 1.9933 | CAG | 2.0254 | GAG | 1.6834 | TAG | 1.2903 |
| AAT | 2.4910 | CAT | 1.8364 | GAT | 1.3358 | TAT | 2.0626 |
| ACA | 2.0131 | CCA | 1.8426 | GCA | 1.4395 | TCA | 1.9583 |
| ACC | 1.1622 | CCC | 1.3141 | GCC | 1.1901 | TCC | 1.5432 |
| ACG | 0.2510 | CCG | 0.2761 | GCG | 0.2381 | TCG | 0.2209 |
| ACT | 1.6076 | CCT | 1.7769 | GCT | 1.3983 | TCT | 2.2134 |
| AGA | 2.2100 | CGA | 0.2205 | GGA | 1.5441 | TGA | 1.9589 |
| AGC | 1.3978 | CGC | 0.2379 | GGC | 1.1896 | TGC | 1.4408 |
| AGG | 1.7749 | CGG | 0.2761 | GGG | 1.3154 | TGG | 1.8462 |
| AGT | 1.6097 | CGT | 0.2516 | GGT | 1.1636 | TGT | 2.0205 |
| ATA | 2.0605 | CTA | 1.2886 | GTA | 1.1347 | TTA | 2.0814 |
| ATC | 1.3343 | CTC | 1.6834 | GTC | 0.9452 | TTC | 1.9728 |
| ATG | 1.8365 | CTG | 2.0269 | GTG | 1.5049 | TTG | 1.8981 |
| ATT | 2.4945 | CTT | 1.9972 | GTT | 1.4607 | TTT | 3.8502 |

**Table 4.** Site-specific haploid mutation rates according to trinucleotide context of base change.

| Site | Site-specific<br>mutation rate<br>($\times 10^{-9}$ /year) | Site | Site-specific<br>mutation rate<br>($\times 10^{-9}$ /year) |
| --- | --- | --- | --- |
| A[C>A]A or T[G>T]T | 1.33 | A[T>A]A or T[A>T]T | 0.43 |
| A[C>A]C or T[G>T]G | 0.73 | A[T>A]C or T[A>T]G | 0.82 |
| A[C>A]G or T[G>T]C | 0.32 | A[T>A]G or T[A>T]C | 0.62 |
| A[C>A]T or T[G>T]A | 0.68 | A[T>A]T or T[A>T]A | 0.40 |
| C[C>A]A or G[G>T]T | 1.89 | C[T>A]A or G[A>T]T | 0.42 |
| C[C>A]C or G[G>T]G | 0.81 | C[T>A]C or G[A>T]G | 0.38 |
| C[C>A]G or G[G>T]C | 0.44 | C[T>A]G or G[A>T]C | 0.61 |
| C[C>A]T or G[G>T]A | 1.05 | C[T>A]T or G[A>T]A | 0.39 |
| G[C>A]A or C[G>T]T | 1.91 | G[T>A]A or C[A>T]T | 0.31 |
| G[C>A]C or C[G>T]G | 1.04 | G[T>A]C or C[A>T]G | 0.29 |
| G[C>A]G or C[G>T]C | 0.85 | G[T>A]G or C[A>T]C | 0.31 |
| G[C>A]T or C[G>T]A | 1.32 | G[T>A]T or C[A>T]A | 0.35 |
| T[C>A]A or A[G>T]T | 0.59 | T[T>A]A or A[A>T]T | 0.26 |
| T[C>A]C or A[G>T]G | 0.50 | T[T>A]C or A[A>T]G | 0.27 |
| T[C>A]G or A[G>T]C | 0.25 | T[T>A]G or A[A>T]C | 0.25 |
| T[C>A]T or A[G>T]A | 0.50 | T[T>A]T or A[A>T]A | 0.25 |
| A[C>G]A or T[G>C]T | 1.09 | A[T>C]A or T[A>G]T | 1.47 |
| A[C>G]C or T[G>C]G | 0.37 | A[T>C]C or T[A>G]G | 0.84 |
| A[C>G]G or T[G>C]C | 0.28 | A[T>C]G or T[A>G]C | 1.84 |
| A[C>G]T or T[G>C]A | 0.65 | A[T>C]T or T[A>G]A | 1.47 |
| C[C>G]A or G[G>C]T | 0.57 | C[T>C]A or G[A>G]T | 0.87 |
| C[C>G]C or G[G>C]G | 0.46 | C[T>C]C or G[A>G]G | 0.81 |
| C[C>G]G or G[G>C]C | 0.25 | C[T>C]G or G[A>G]C | 1.10 |
| C[C>G]T or G[G>C]A | 0.53 | C[T>C]T or G[A>G]A | 0.74 |
| G[C>G]A or C[G>C]T | 0.92 | G[T>C]A or C[A>G]T | 0.67 |
| G[C>G]C or C[G>C]G | 0.66 | G[T>C]C or C[A>G]G | 0.56 |
| G[C>G]G or C[G>C]C | 0.63 | G[T>C]G or C[A>G]C | 0.65 |
| G[C>G]T or C[G>C]A | 0.76 | G[T>C]T or C[A>G]A | 0.95 |
| T[C>G]A or A[G>C]T | 0.50 | T[T>C]A or A[A>G]T | 0.57 |
| T[C>G]C or A[G>C]G | 0.74 | T[T>C]C or A[A>G]G | 0.54 |
| T[C>G]G or A[G>C]C | 0.55 | T[T>C]G or A[A>G]C | 0.62 |
| T[C>G]T or A[G>C]A | 0.85 | T[T>C]T or A[A>G]A | 0.50 |
| A[C>T]A or T[G>A]T | 3.21 | A[T>G]A or T[A>C]T | 0.20 |
| A[C>T]C or T[G>A]G | 2.46 | A[T>G]C or T[A>C]G | 0.16 |
| A[C>T]G or T[G>A]C | 9.72 | A[T>G]G or T[A>C]C | 0.44 |
| A[C>T]T or T[G>A]A | 2.80 | A[T>G]T or T[A>C]A | 0.20 |
| C[C>T]A or G[G>A]T | 2.41 | C[T>G]A or G[A>C]T | 0.16 |
| C[C>T]C or G[G>A]G | 2.65 | C[T>G]C or G[A>C]G | 0.19 |
| C[C>T]G or G[G>A]C | 5.34 | C[T>G]G or G[A>C]C | 0.41 |
| C[C>T]T or G[G>A]A | 4.69 | C[T>G]T or G[A>C]A | 0.25 |
| G[C>T]A or C[G>A]T | 3.96 | G[T>G]A or C[A>C]T | 0.14 |
| G[C>T]C or C[G>A]G | 5.60 | G[T>G]C or C[A>C]G | 0.08 |
| G[C>T]G or C[G>A]C | 18.84 | G[T>G]G or C[A>C]C | 0.26 |
| G[C>T]T or C[G>A]A | 7.07 | G[T>G]T or C[A>C]A | 0.15 |
| T[C>T]A or A[G>A]T | 1.09 | T[T>G]A or A[A>C]T | 0.14 |
| T[C>T]C or A[G>A]G | 1.45 | T[T>G]C or A[A>C]G | 0.16 |
| T[C>T]G or A[G>A]C | 3.28 | T[T>G]G or A[A>C]C | 0.32 |
| T[C>T]T or A[G>A]A | 1.40 | T[T>G]T or A[A>C]A | 0.19 |

Although for convenience we enumerate the haploid mutation rates for all 192 site contexts (Supplementary Table 4) it is important to note that the rates at a particular site context and its complementary partner, e.g. A[C>A]A and T[G>T]T, cannot be distinguished since only the sum of their rates is measured. Thus, in Supplementary Table 4, the first entry 1.33 *×* 10^−9^ is the sum of the rates of both A[C>A]A and T[G>T]T mutations, which is the relevant rate for calculating how frequently a site mutates since either strand could have undergone the mutation.

The size of each study’s panel ‘footprint’ was determined from the study’s published information. Study-specific mutation rates were then calculated using a custom Python script for each gene or variant of interest, by summing the site-specific mutation rates for the regions at which the study called variants (Tables 5 and 6 and Supplementary Figures 3, 4, 5 and 6).

**Table 5.**
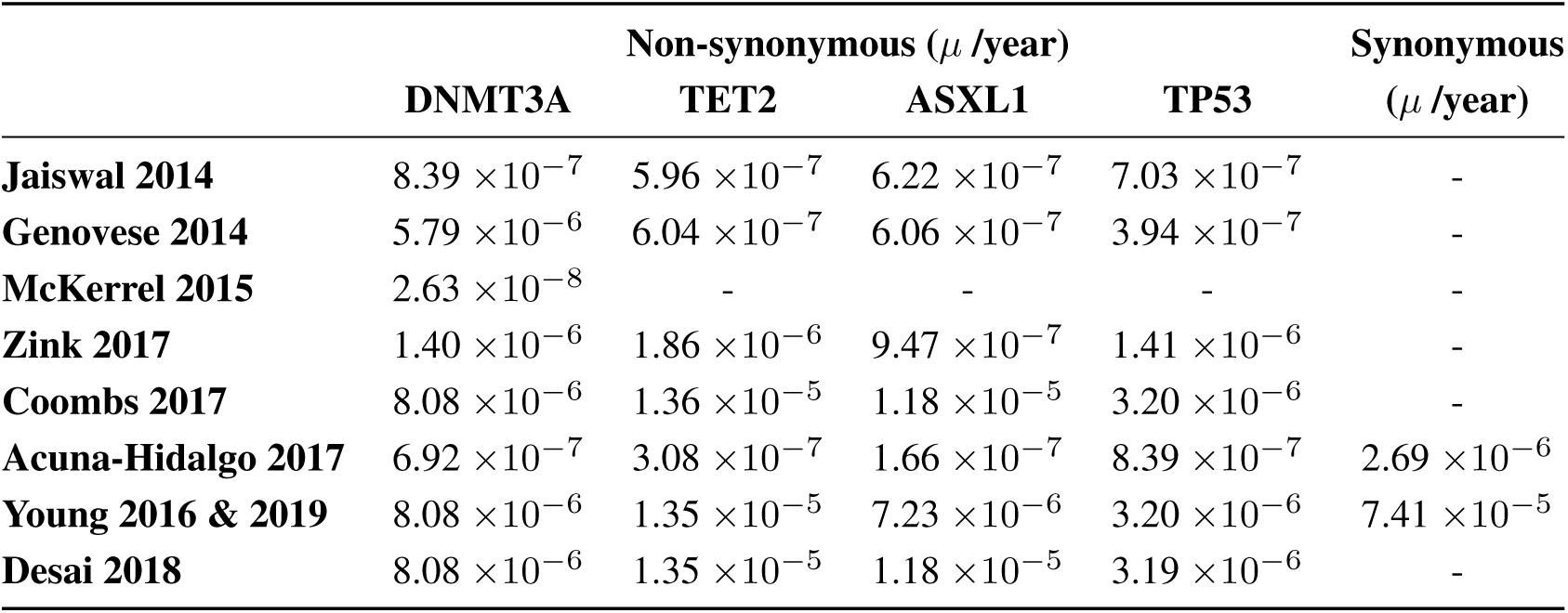
Study-specific mutation rates for non-synonymous DNMT3A, TET2, ASXL1, TP53 variants and all synonymous variants. Calculated by summing the site-specific mutation rates (Supplementary Table 4) across the regions of the gene covered by each study.

**Table 6.** Variant-specific mutation rates for the top 20 most commonly observed variants. Calculated using the site-specific mutation rates (Supplementary Table 4) for the nucleotide change (and its trinucleotide context) that gives rise to the variant.

| Variant | Trinucleotide Context | $\mu$ /year |
| --- | --- | --- |
| **DNMT3A R320\*** | C[C>T]G | $5.34 \times 10^{-9}$ |
| **DNMT3A R326C** | C[C>T]G | $5.34 \times 10^{-9}$ |
| **DNMT3A R598\*** | G[C>T]G | $1.88 \times 10^{-8}$ |
| **DNMT3A R729W** | C[C>T]G | $5.34 \times 10^{-9}$ |
| **DNMT3A Y735C** | T[A>G]C | $1.84 \times 10^{-9}$ |
| **DNMT3A R736C** | C[C>T]G | $5.34 \times 10^{-9}$ |
| **DNMT3A R736H** | C[G>A]C | $1.88 \times 10^{-8}$ |
| **DNMT3A R771\*** | G[C>T]G | $1.88 \times 10^{-8}$ |
| **DNMT3A R882C** | C[C>T]G | $5.34 \times 10^{-9}$ |
| **DNMT3A R882H** | C[G>A]C | $1.88 \times 10^{-8}$ |
| **DNMT3A W860R** | A[T>A]G, A[T>C]G | $2.46 \times 10^{-9}$ |
| **DNMT3A P904L** | C[C>T]G | $5.34 \times 10^{-9}$ |
| **GNB1 K57E** | C[A>G]A | $9.53 \times 10^{-10}$ |
| **IDH2 R140Q** | C[G>A]G | $5.60 \times 10^{-9}$ |
| **JAK2 V617F** | T[G>T]T | $1.33 \times 10^{-9}$ |
| **SF3B1 K666N** | A[G>T]A, A[G>C]A | $1.35 \times 10^{-9}$ |
| **SF3B1 K700E** | G[A>G]A | $7.41 \times 10^{-10}$ |
| **SRSF2 P95H** | C[C>A]C | $8.15 \times 10^{-10}$ |
| **SRSF2 P95L** | C[C>T]C | $2.65 \times 10^{-9}$ |
| **SRSF2 P95R** | C[C>G]C | $4.56 \times 10^{-10}$ |

**Supplementary Figure 3.**
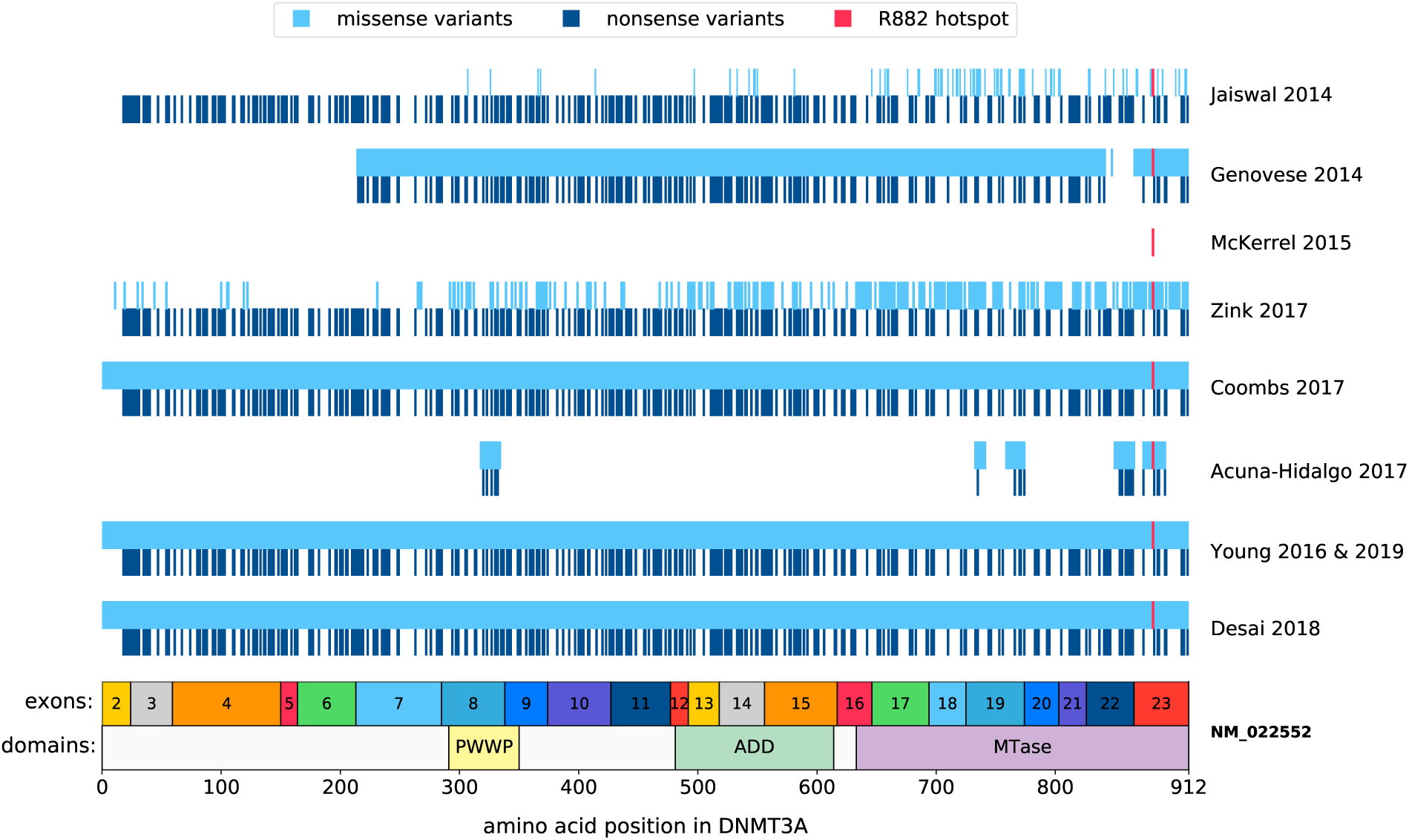
Panel ‘footprint’ for DNMT3A for each of the studies included in our analysis. Regions were inferred from each study’s published information. PWWP: Pro-Trp-Trp-Pro domain, ADD: ATRX-DNMT3A-DNMT3L domain, MTase: Methyltransferase domain.

**Supplementary Figure 4.**
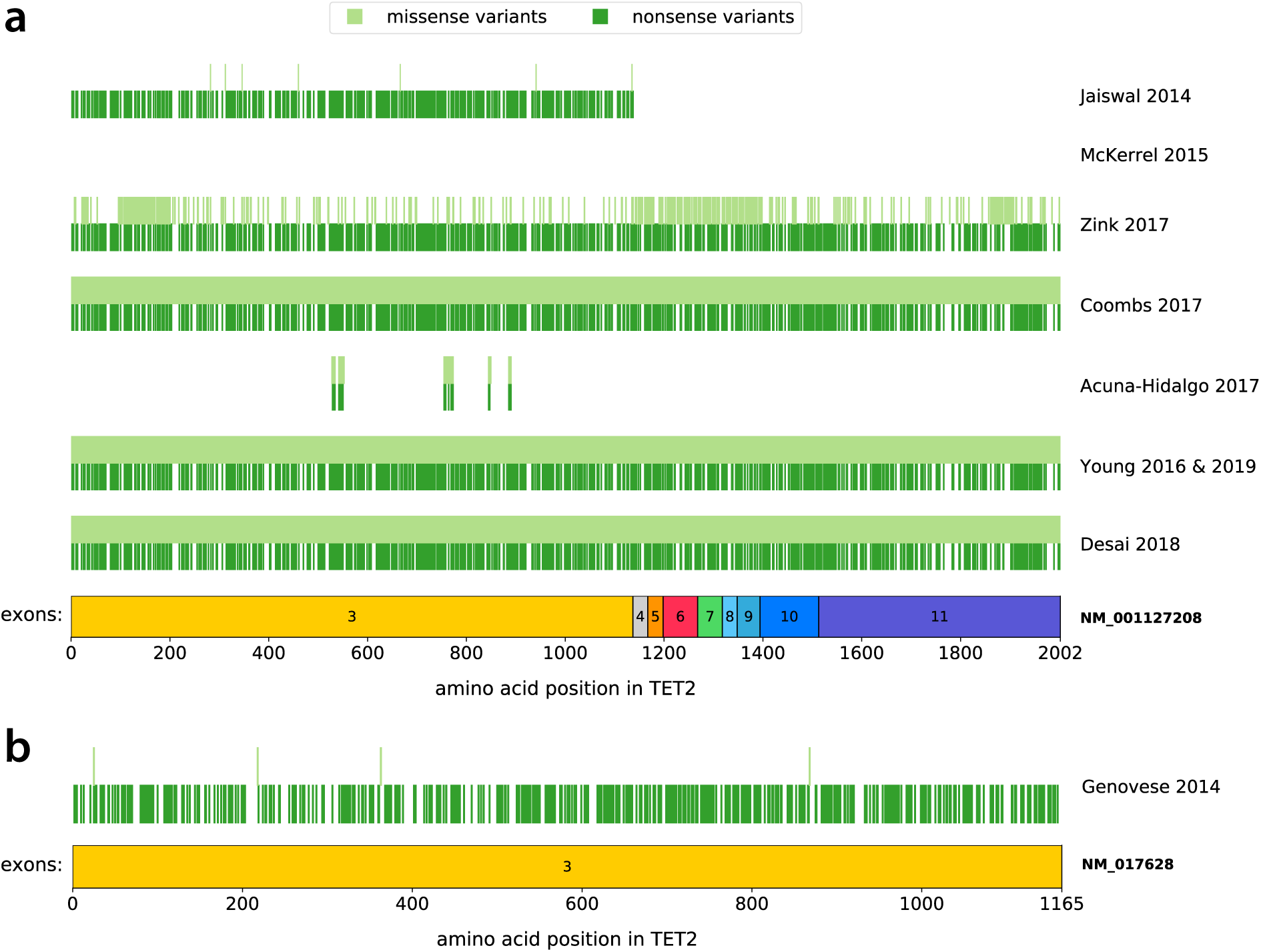
Panel ‘footprint’ for TET2 for each of the studies included in our analysis. Regions were inferred from each study’s published information. **a:** The majority of studies annotated variants using the NM_001127208 transcript. **b:** The NM_017628 transcript was used by Genovese 2014. McKerrel 2015 did not target TET2 in their panel.

**Supplementary Figure 5.**
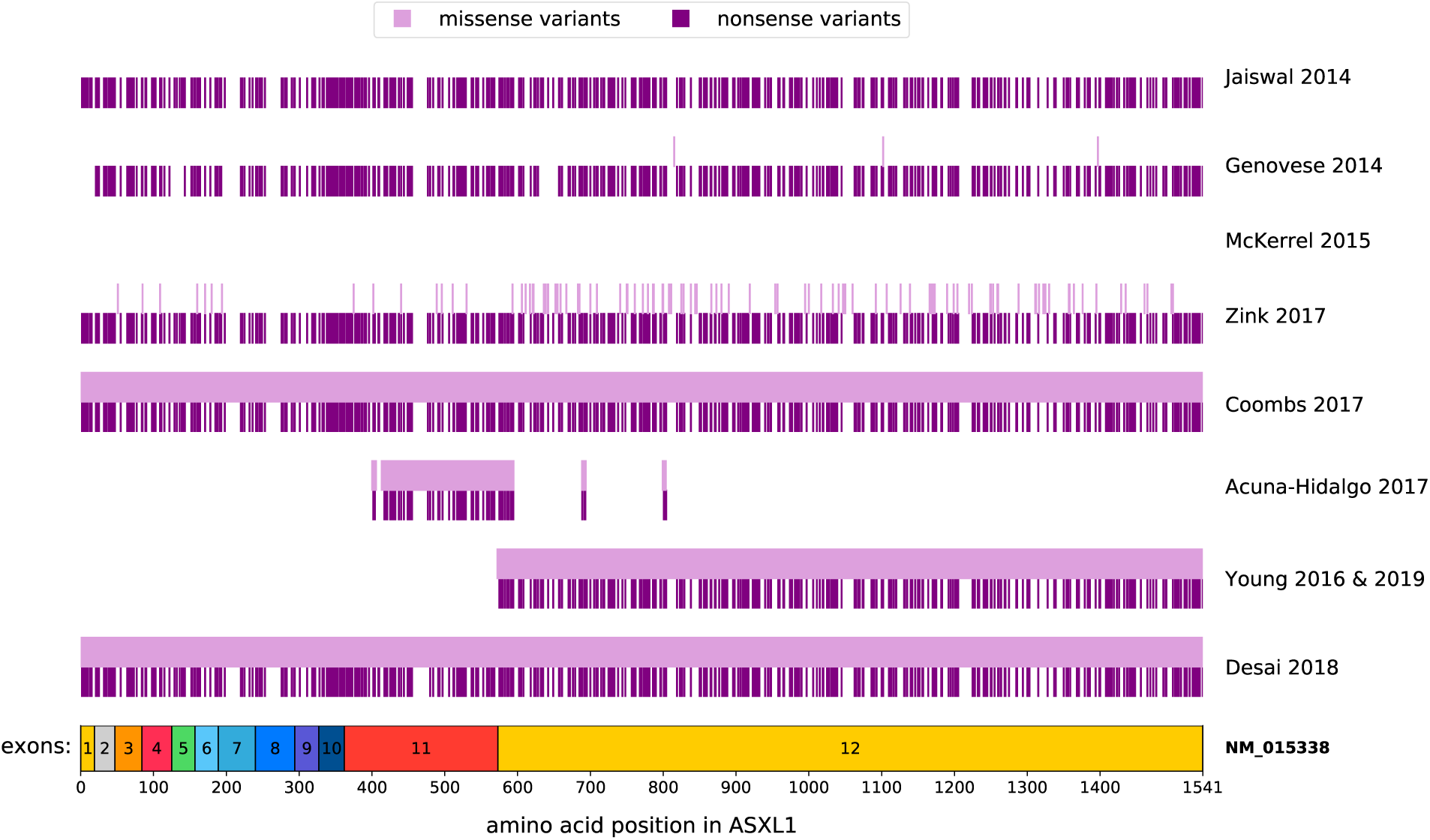
Panel ‘footprint’ for ASXL1 for each of the studies included in our analysis. Regions were inferred from each study’s published information. The majority of studies annotated variants using the NM_015338 transcript. McKerrel 2015 did not target ASXL1 in their panel.

**Supplementary Figure 6.**
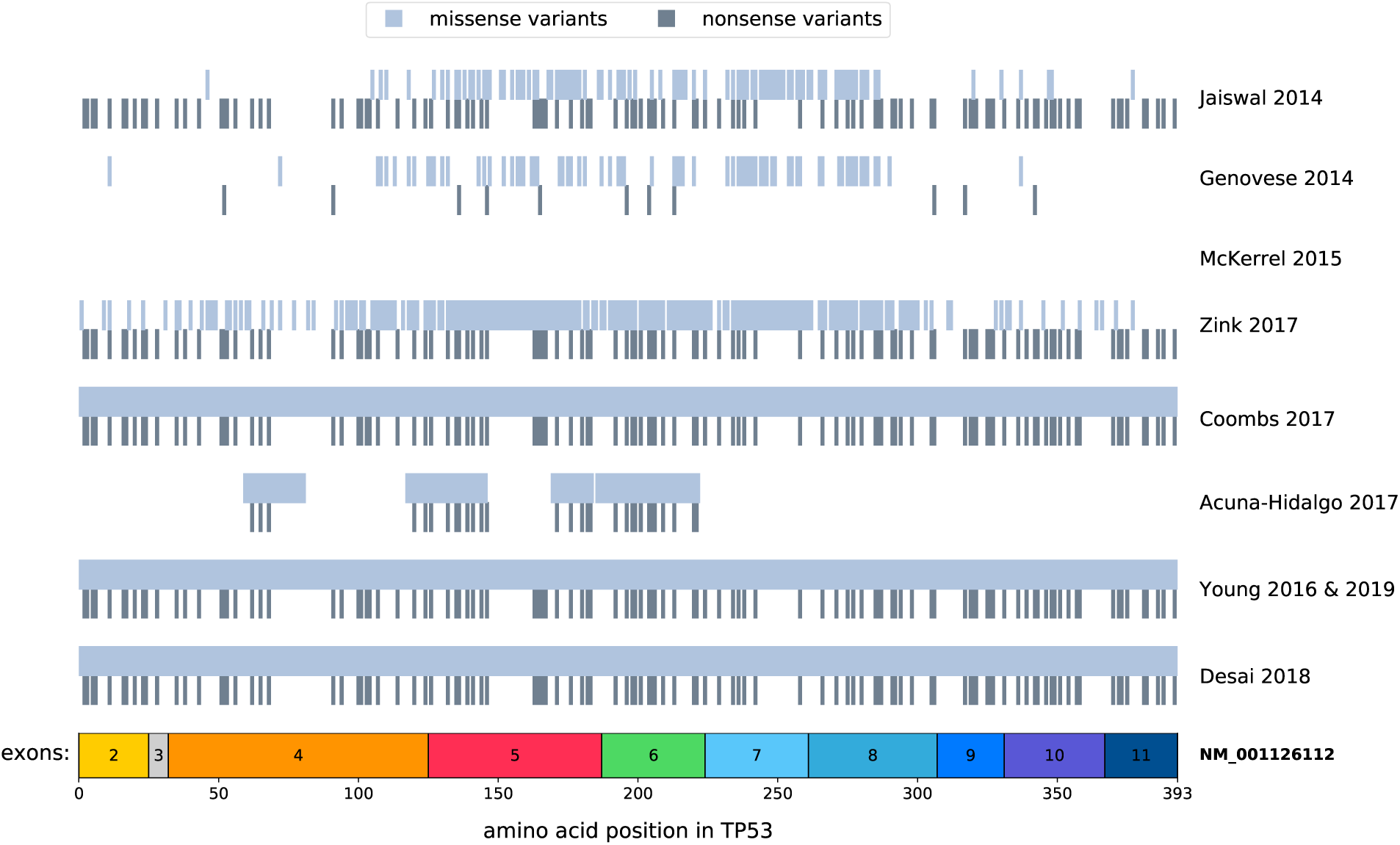
Panel ‘footprint’ for TP53 for each of the studies included in our analysis. Regions were inferred from each study’s published information. The majority of studies annotated variants using the NM_001126112 transcript. McKerrel 2015 did not target TP53 in their panel.

## Supplementary Note 5: The emergence of clones with multiple drivers

Clones harbouring multiple driver mutations are expected to be rare, even at late ages. The analysis depends on the spectrum of fitness effects, but the main principle can be understood more easily by considering highly fit single-mutants that are acquired at rate *U*_*b*_ and expand at rate *s*, and which themselves can acquire a second driver mutation at rate *U*_*b*_ which confers an additional fitness effect to the cell, which, in combination with the first mutation, confers a total fitness of *αs*, to the cell. In this case, the double mutant clone enters the HSC population at a rate 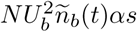. It then grows exponentially, but at a rate *α* times that of the first clone, where *α >* 1. The clone size distribution of clones with two driver mutations that results from this is

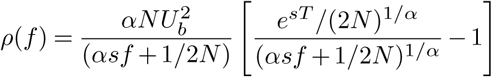

for time *T > t*.

On a log-log density plot, this distribution has a predominantly slope of – 1*/α* due to the effective ∝ *f* ^−(1+1*/α*)^ relationship. This is valid for all positive *α* and it is clear that the hitchhikers are a special case of this scenario with *α* = 1. Because the mutation rates *U*_*b*_ are low, the expected number of clones with multiple drivers is also low over the timescales of a human lifespan. For example, the number of expected double-mutants detected at VAFs>0.5% in the case where the second mutant confers the same additional advantage as the first (*α* = 2) with *s* ≈ 10% and *U*_*b*_ = 10^−5^ in an HSC population of 10^5^, is 0.03% at age 40, 0.47% at age 60, and 3.75% at age 80. Clones with more than two drivers mutations are even rarer.

Extending this to predict the prevalence of double-mutant clones from within ten of the most commonly mutated CH genes and taking in to account the distribution of fitness effects across these genes (Supplementary Note 9), we can predict that, at a VAF detection limit of 1%, <10% of individuals aged 100 will harbour two mutations from within these genes within the same clone (Supplementary Figure 7).

**Supplementary Figure 7.**
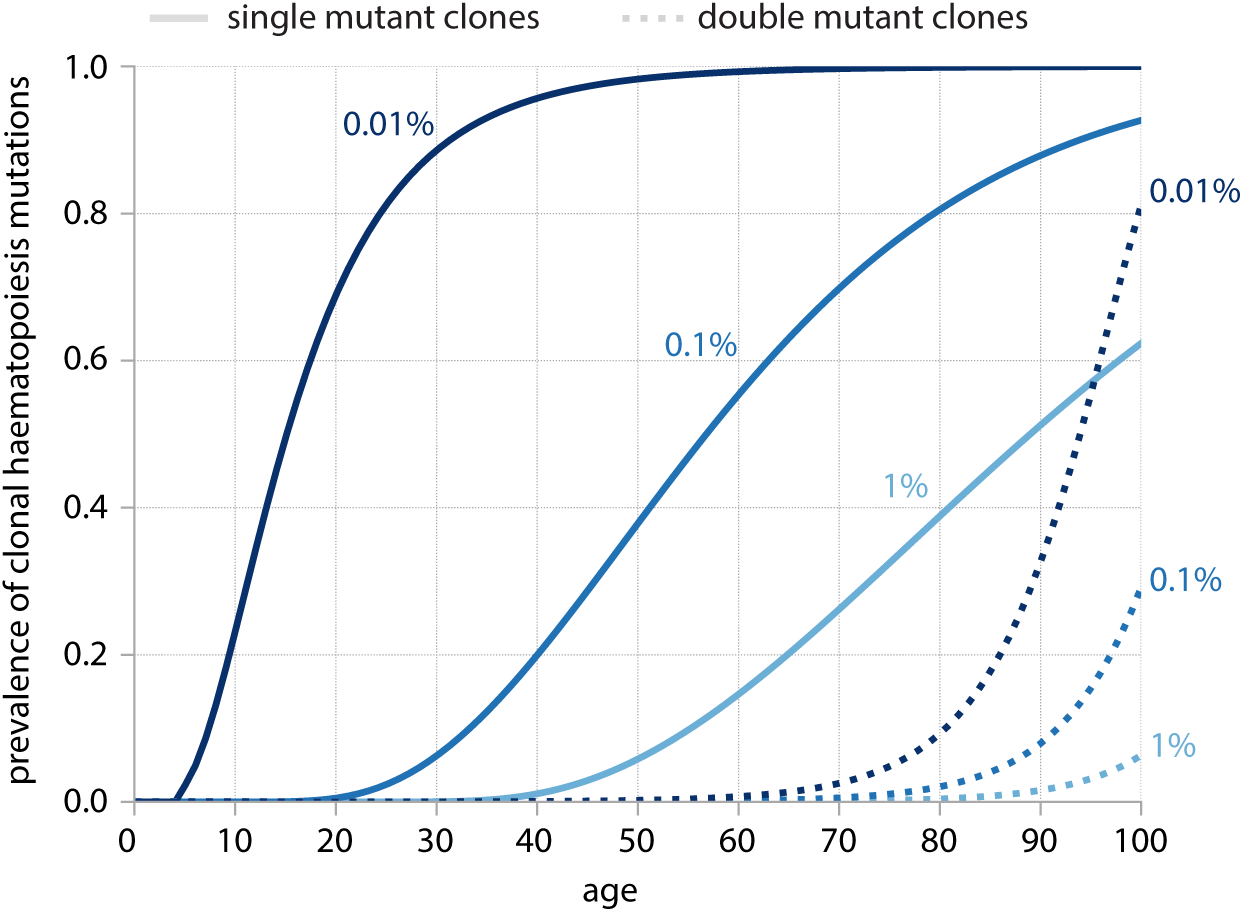
Prevalence of single-mutant clones and double-mutant clones as function of age for different sequencing thresholds. Prevalence is predicted for variants to have been acquired within 10 of the most commonly mutated CH genes (DNMT3A, TET2, ASXL1, JAK2, TP53, CBL, SF3B1, SRSF2, IDH2 and KRAS), taking in to account the distribution of fitness affects across these genes (Supplementary Note 9). Single-mutant clones (clones harbouring one variant from within these 10 genes) are represented by the solid lines and double-mutant clones (clones harbouring two variants from within these 10 genes) are represented by dashed lines. Lines are colour-coordinated according to VAF limit of detection (0.01%, 0.1% or 1%).

## Supplementary Note 6: Parameter estimation using Maximum Likelihood Estimations

Parameter estimation was performed using a custom Python script. Briefly, probability density histograms, as a function of log VAFs, were plotted, using Doane’s method for log VAF bin size calculation ^58^. To enable comparison between studies, the densities were normalised by dividing by [number of individuals in the study × bin widths] and, in order to read *Nτ* from the y-intercept, the densities were then rescaled by dividing by 2*µ*, where *µ* is the study-specific haploid mutation rate for the region (or variant) being plotted (Supplementary Note 2, Tables 5 and 6).

### Parameter estimation for top 20 most commonly observed variants

Estimates for *Nτ* and *s* for DNMT3A R882H variants were inferred first, using a maximum likelihood approach. We minimised the L2 norm between the log rescaled densities and the predicted densities, for all data points, obtained by integrating the theoretical density for a given age (eqn. 9) with a distribution of ages (normal distribution with mean 55 years and standard deviation *σ*) and optimised *σ* along with *Nτ* and *s* (Supplementary Figure 8). The optimal *σ* was 11.4 years, broadly consistent with the age range reported in most studies, with *Nτ ≈* 100,000 years and *s ≈* 15% (Supplementary Figure 8). Because our estimate for *Nτ* agrees with other independent estimates,^36^ to calculate the *s* for other individual variants we fixed *Nτ* and *σ* to that inferred from R882H (*≈* 100,000 years) and used a maximum likelihood approach to optimise *s* as well as *µ* (Supplementary Figure 9). Data points ≥ 50% VAF were not included in maximum likelihood analyses. Of note, in most cases the inferred value of *µ* agreed to within a factor of 3 to that estimated by the site-specific trinucleotide context (Supplementary Table 6).

### Relative fitness effects of DNMT3A R882H and DNMT3A R882C

A key feature of our framework is its ability to disentangle the relative effects of mutation rate and fitness effect. For example, R882H (*n* =105) is the most commonly observed variant in DNMT3A, across all nine studies, followed by R882C (*n* =61), but is R882H’s high prevalence explainable by a higher mutation rate (1.88 × 10^−8^ vs 5.34 × 10^−9^ for R882H and R882C respectively), higher fitness effect or both? Plotting the distribution of VAFs for each of these variants and using maximum likelihood approaches, as above, to infer fitness effects, reveals that R882C actually has a higher fitness effect than R882H (Supplementary Figure 10). So, although R882H variants are observed most frequently, this is attributable to a high mutation rate in combination with a high fitness and R882C mutations actually have a higher fitness effect and are thus potentially more pathogenic.

**Supplementary Figure 8.**
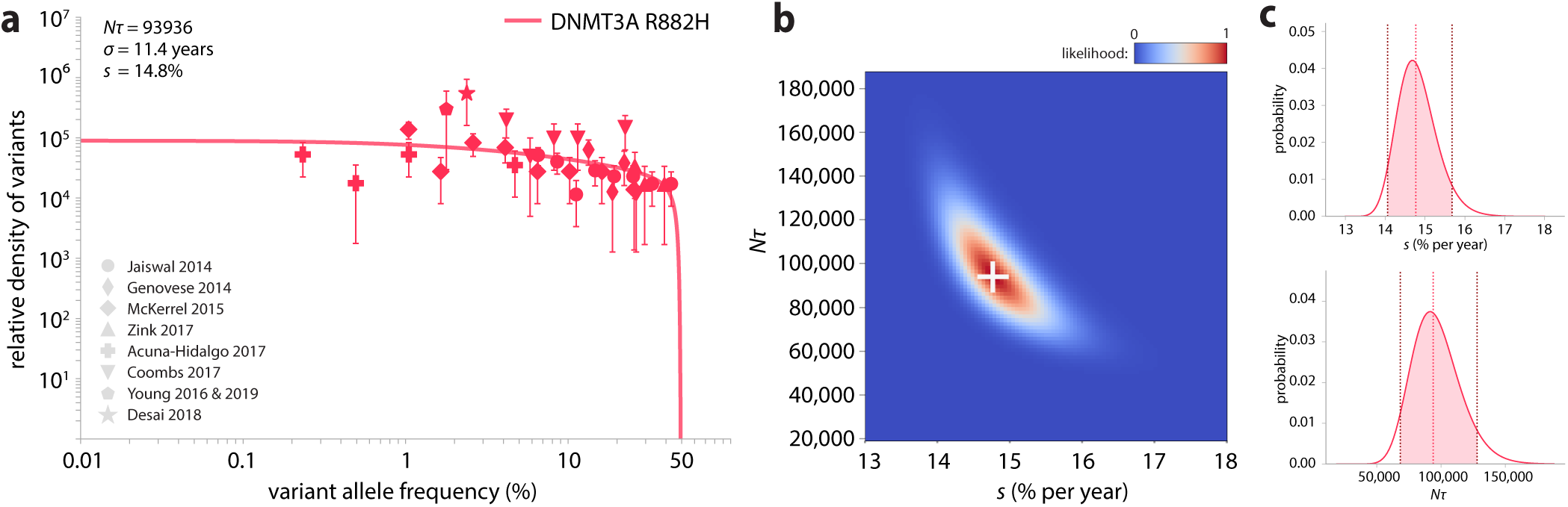
Parameter estimation for DNMT3A R882H. (a) Probability density histogram for R882H with theory distribution fitted using maximum likelihood estimates. The mean age was fixed at 55 years (normally distributed) and maximum likelihood approaches were used to infer the standard deviation of ages (*σ*) = 11.4 years, *Nτ* = 93936 and *s* = 14.8% per year. **(b) Maximum likelihood heatmap for** *Nτ* **and** *s* **estimates for R882H.** White cross marks the most likely *Nτ* (93936) and *s* (14.8% per year). **(b) Distribution of likelihoods for** *s* **and** *Nτ*. Red vertical line represents most likely value. 95% confidence intervals are shown shaded in pink: *Nτ* 68317 – 128094, *s* 14.0 – 15.7%.

**Supplementary Figure 9.**
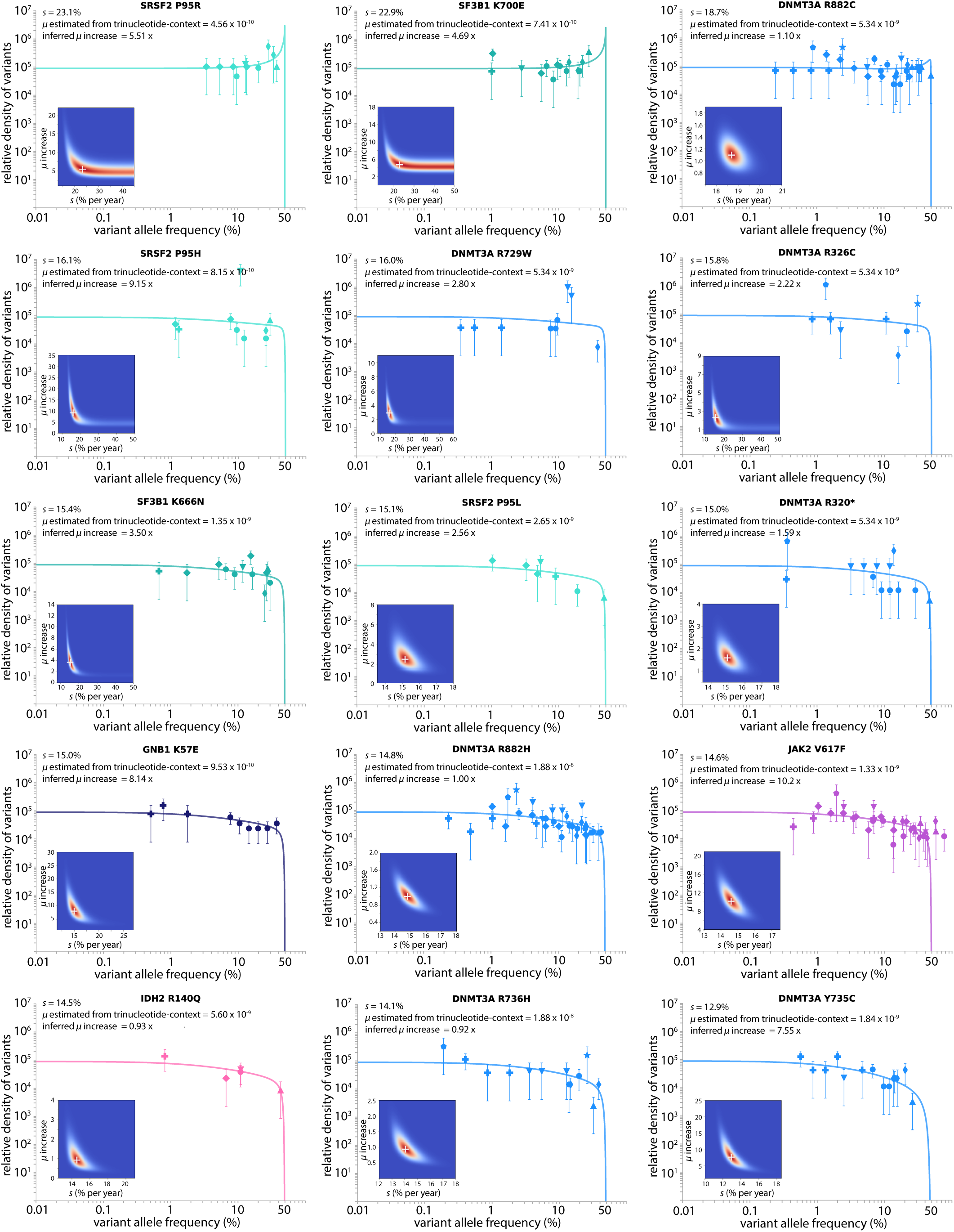
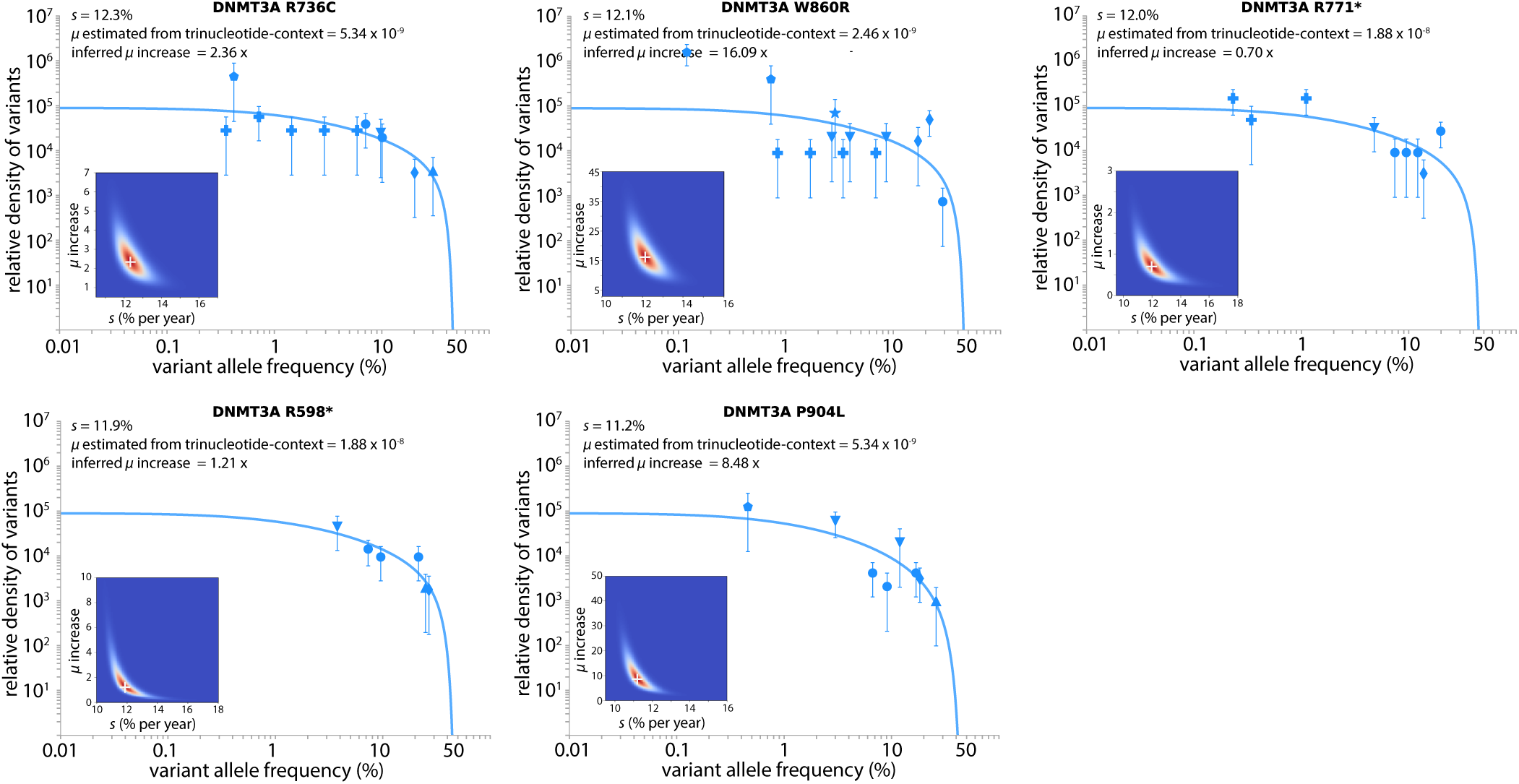
Parameter estimation for the top 20 most commonly observed CH variants. *Nτ* and the standard deviation of ages (*σ*) were fixed to that inferred from DNMT3A R882H and maximum likelihood approaches were used to infer *s* for each variants, as well as the increase of *µ* relative to the *µ* estimated from the variant’s site-specific trinucleotide context (Supplementary Table 6). Each study is represented by a shaped symbol as described in Figure 1e.

**Supplementary Figure 10.**
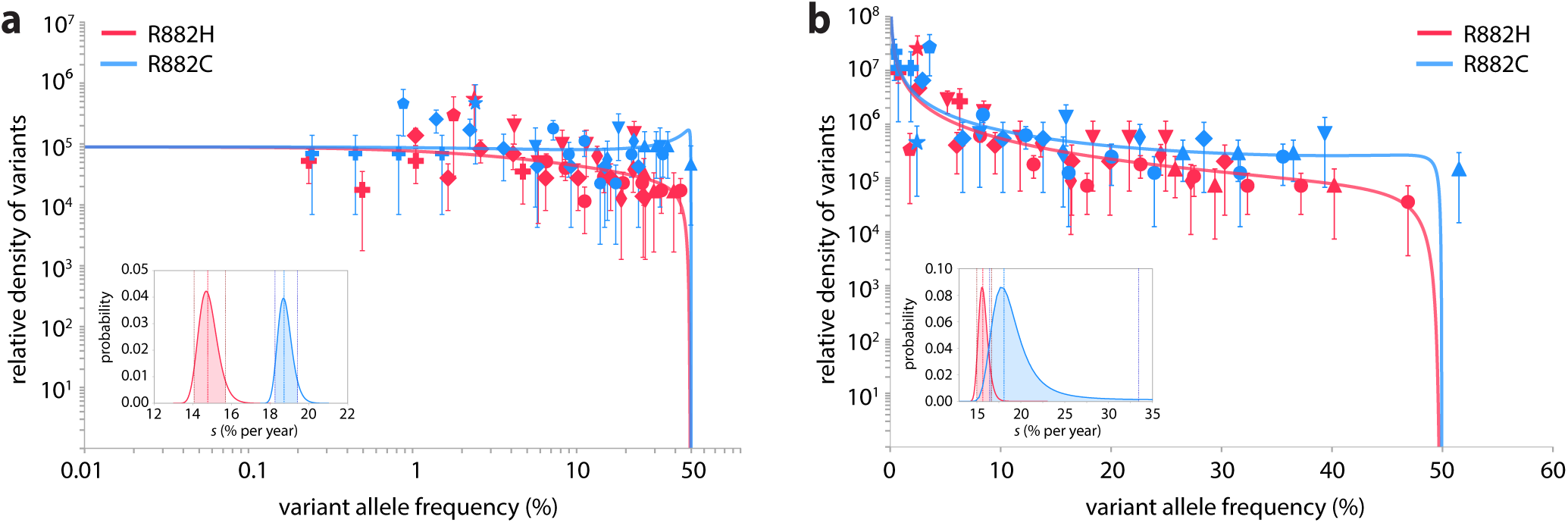
Fitted theory distributions for DNMT3A R882H and R882C using Maximum Likelihood Estimates. **(a) Log-log plot.** Probability density histograms for R882H (red data) and R882C (blue data), as a function of log VAFs, plotted using Doane’s method for logarithmic VAF bin size calculation. *Nτ* was fixed to ≈ 100,000 and maximum likelihood approaches were used to infer *s* of 14.8% (95%C.I. 14.1-15.7%, pink shaded area) for R882H and *s* of 18.7% (95% C.I.18.2-19.4%, blue shaded area) for R882C. **(b) Linear-log plot.** Probability density histograms for R882H (red data) and R882C (blue data), as a function of linear VAFs, plotted using Doane’s method for linear VAF bin size calculation. *Nτ* was fixed to ≈ 100,000 and maximum likelihood approaches were used to infer *s* of 15.7% (95% C.I. 14.9-16.6 %, pink shaded area) for R882H and *s* of 18.2% (95% C.I. 16.5-33.4%, blue shaded area) for R882C. Each study is represented by a shaped symbol as described in Figure 1e.

**Supplementary Figure 11.**
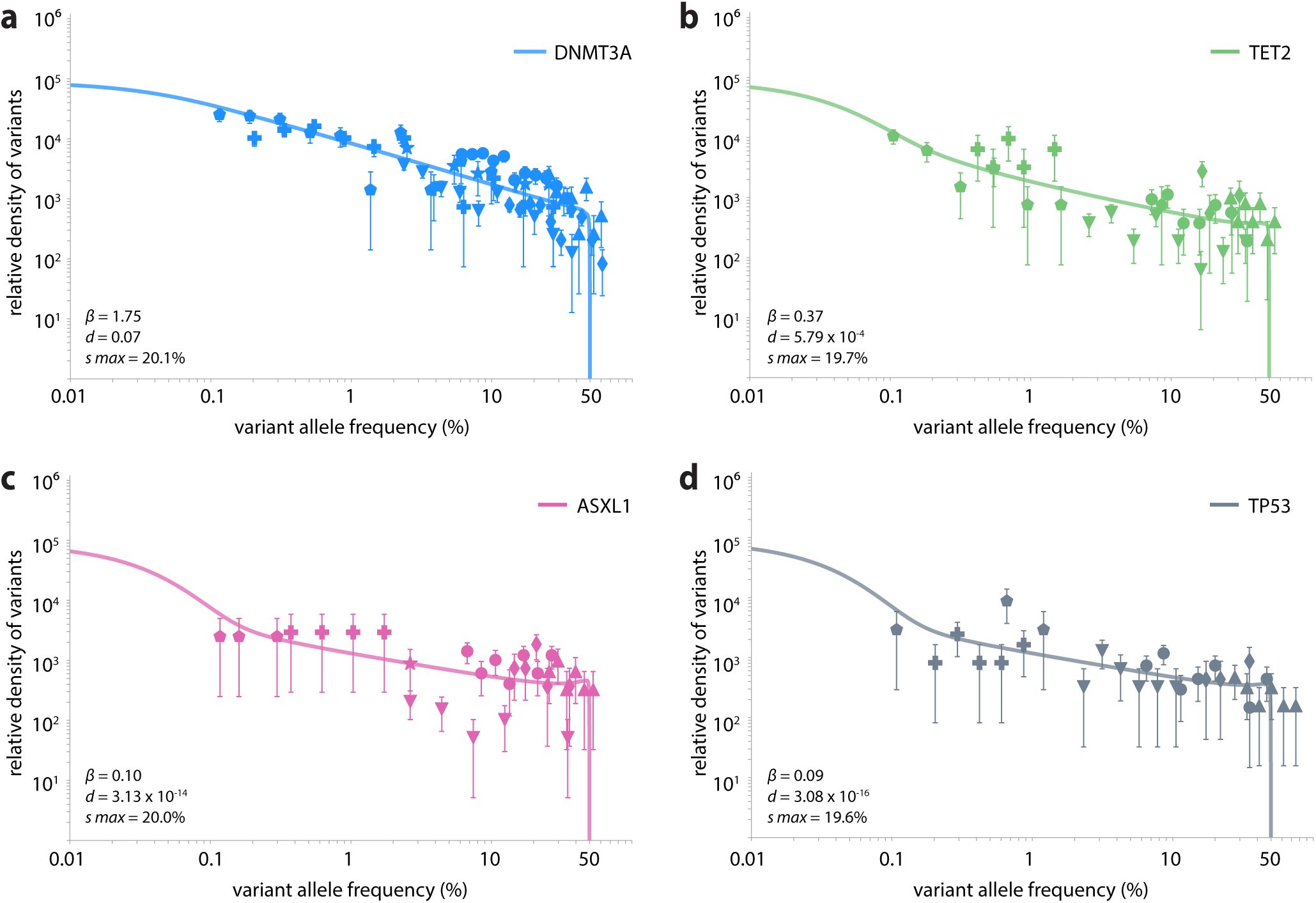
Parameter estimation for distribution of fitness effects of nonsynonymous variants within commonly mutated CH genes. **(a)** DNMT3A. **(b)** TET2. **(c)** ASXL1. **(d)** TP53. Each study is represented by a shaped symbol as described in Figure 1e.

### Parameter estimation for distribution of fitness effects within genes: DNMT3A, TET2, ASXL1, TP53

For nonsynonymous variants within DNMT3A, TET2, ASXL1 and TP53, estimates for the distribution of *s* were inferred by fixing *Nτ* and the standard deviation of ages (*σ*) to that inferred from DNMT3A R882H. We parameterised the distribution of fitness effects using a family of stretched exponential distributions,

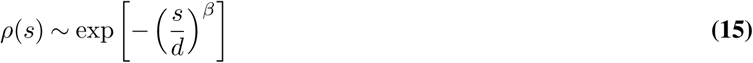

with a maximum *s* = *s*_*max*_. We then performed a maximum likelihood procedure, optimising the shape (*β*) and scale (*d*) of the distribution as well *s*_*max*_ (Supplementary Figure 11).

### Parameter estimation for synonymous variants

Only three studies (Young 2016^10^, Young 2019^15^ and Acuna-Hidalgo 2017^12^) reported synonymous variants. Acuna-Hidalgo’s study included 2006 participants with a uniform distribution of ages. Since Young 2016 and 2019 contained a total of 89 participants, which is small relative to Acuna-Hidalgo, we assumed a uniform distribution of ages across all three studies combined. The maximum likelihood estimations for synonymous variants were therefore calculated over a uniform distribution of ages, from 20-69 years old. Fixing *Nτ* to that inferred from DNMT3A R882H, maximum likelihood approaches inferred a *ϕ* value of 0.11%, ∼4.5-fold higher than the predicted *ϕ* value for neutral variants (0.024%) (Supplementary Figure 12a). But, if the synonymous variants >0.25% VAF were assumed to be potential hitchhiker mutations and the maximum likelihood analysis included only synonymous variants <0.25% VAF, the inferred *ϕ* was 0.03%, which is within a factor of 1.3 of the value predicted for neutral variants (Supplementary Figure 12b).

**Supplementary Figure 12.**
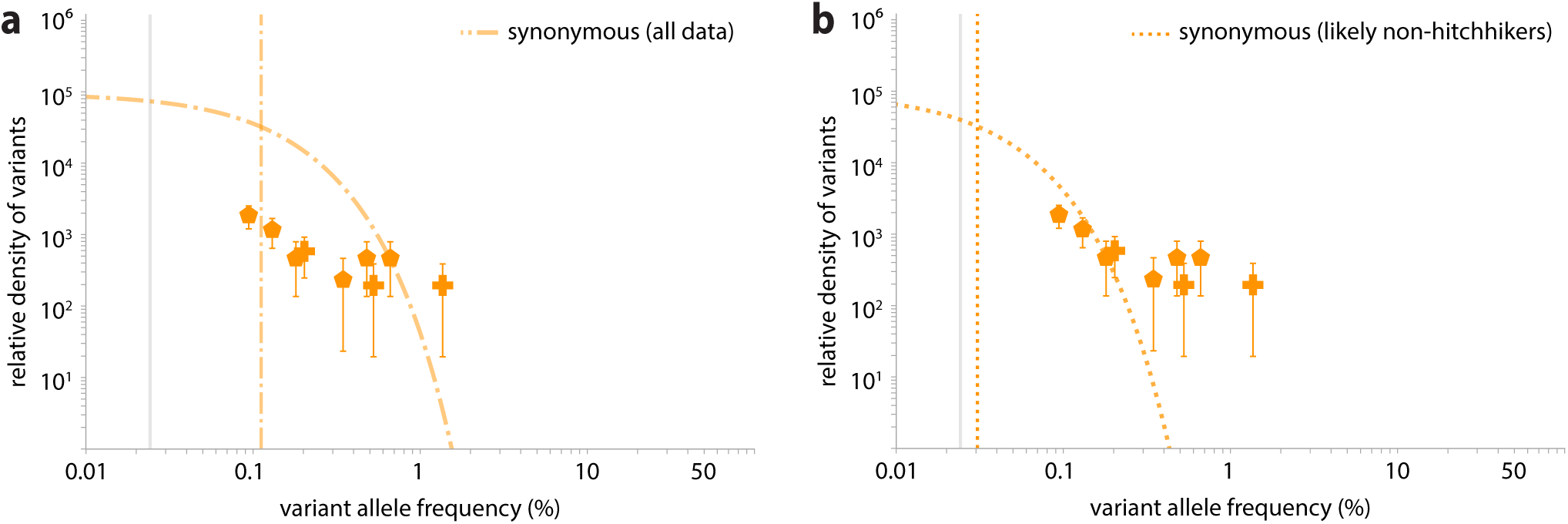
Fitted theory distribution for synonymous variants. **(a)** If synonymous variants of any VAF are included in the maximum likelihood approach, the inferred *ϕ* value (orange dot-dashed line) is 4.6-fold higher than the predicted *ϕ* value (grey vertical line). **(b)** If synonymous variants >0.25% VAF are assumed to be hitchhikers, maximum likelihood approaches on those <0.25% VAF infer a *ϕ* value (orange dashed line) only 1.3-fold higher than the predicted *ϕ* value (grey vertical line). Each study is represented by a shaped symbol as described in Figure 1e.

## Supplementary Note 7: Developmental mutations and neutral ‘hitchhikers’

### Developmental mutations

Mutations occurring during early development, while the number of HSCs expands, can be present in high frequencies in the adult HSC population. Assuming the entire HSC population in a person grows as *e*^*rt*^ from a single ancestor, the probability a neutral mutation enters at time t is *ρ*(*t*) = *U*_*n*_*e*^*rt*^, where *U*_*n*_ is the neutral mutation rate per cell division. Under the deterministic assumption (only used during the development phase), the neutral mutant lineage that entered grows as *f* = (1*/*2*N*)*e*^*r*(*T*^*d*^*-t*)^. The full site frequency spectrum of developmental mutations will therefore be

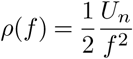

This can be used to estimate how likely it is that high VAF synonymous variants were early developmental mutations. The neutral mutation rate per cell division during development and is tightly constrained by data from previous studies ^36^, where it is found there are an estimated 1.2 mutations per HSC division per genome during development. Thus, for a given study *U*_*n*_ = 1.2 × (*g/G*), where *g* is the number of synonymous bases covered by the sequencing panel used in that study and *G* the genome size. The densities in Figure 1e of the main text have been rescaled by dividing though by twice the study-specific neutral mutation rate per year, which is 2*g* × *u*_*n*_, where *u*_*n*_ *≈* 5 × 10^−9^ is the mutation rate per bp per year. Therefore the amplitude of the developmental variant distribution should be

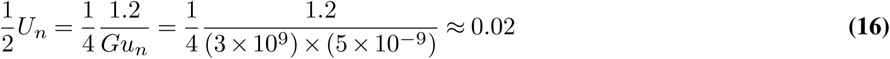

The amplitude of the 1*/f* ^2^ dashed line in Figure 1e is *≈* 1, which is 50-fold larger than expected for developmental mutations. Therefore, it is unlikely the observed high VAF synonymous variants are developmental in origin. In fact, by integrating the density of developmental mutations from each study, we determined that the number of synonymous variants with VAF>0.3% that are developmental in origin to be <1.

### Genetic hitchhikers

An alternative explanation for high-VAF synonymous variants is that they ‘hitchhike’ alongside a beneficial mutation. This can happen in two distinct ways: the neutral mutation occurs inside an expanding beneficial clone, or the neutral clone was lucky enough to survive drift and then acquired a subsequent beneficial mutation. In the first case, a constant wild-type stem cell population feeds beneficial mutations at rate *µ*_*b*_ per year, which results in the total size of beneficial lineages growing as

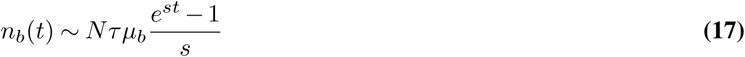

The density of hitchhikers that establish in the interval of time between [*t, t* + *dt*] years is *µ*_*n*_*n*_*b*_(*t*)*s dt*. Since a neutral hitchhiker occurring at time, *t*, will reach a size *n ≈* (*e*^*s*(*T-t*)^ – 1)*/s* by time *T*, the variable *t* can be eliminated to express the density in terms of *f* yielding

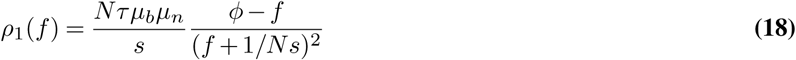

where *ϕ* is the characteristic maximum VAF *ϕ* = (*e*^*st*^ *–* 1)*/Ns* on which the synonymous mutations hitchhike. On a log-log plot, this distribution appears as a straight line with gradient -1 at intermediate frequencies due to the effective ∼ 1*/f* ^2^ scaling, a consequence of exponential growth of clones being fed at exponential rates, analogous to the Luria-Delbruck distribution ^59^. At high frequencies, the density of hitchhikers falls sharply as beneficial clones only grow linearly at very early times, which is when the high-frequency hitchhikers enter the stem cell population. The form of the distribution at intermediate frequencies is the same as for developmental mutations, having a 1*/f* ^2^ scaling law. The amplitude of the 1*/f* ^2^ distribution is approximately *Nτµ*_*b*_*µ*_*n*_*ϕ/s*. In Figure 1e, the density has been rescaled by dividing through by twice the neutral mutation rate 2*µ*_*n*_ hence the amplitude for the density of high VAF synonymous variants should be approximately *Nτµ*_*b*_*ϕ/*2*s*. Using our estimated *Nτ ≈* 50, 000 years and that fact that the ratio *ϕ/*2*s* ∼ 1 for variants with *s ≈* 10% per year at age 60 years and *ϕ/*2*s* ∼ 0.1 for variants with *s ≈* 5% per year at age 60 years, in order to fit the data we require a genome-wide beneficial mutation rate of *µ*_*b*_ = 2 × 10^−5^ per year to fitness effects *s >* 10% and *µ*_*b*_ = 2 × 10^−4^ per year to fitness effects *s >* 5% per year. These numbers are plausible: the inferred mutation rate to fitness effects *s >* 10% per year across ten of the most commonly mutated CH genes alone is *µ*_*b*_ *≈* 1.5 × 10^−6^ per year. While this is 10-fold smaller than required to explain the density of synonymous variants reaching high VAFs, it suggests there may be many more highly fit mutations elsewhere in the genome, outside these 10 commonly mutated CH genes.

In the second case, a hitchhiker occurs by a neutral clone acquiring a beneficial mutation. The constant wild-type stem cell population feeds neutral mutations at rate *µ*_*n*_, which results in the total size of neutral lineages growing as

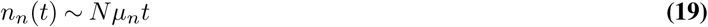

Hitchhikers enter with a probability of approximately *µ*_*b*_*n*_*n*_(*t*)*s* at time *t*. Performing the same change of variables to eliminate *t* in favour of *f* yields

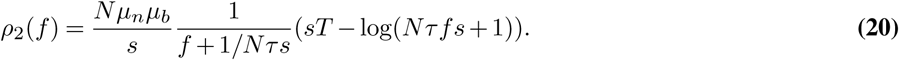

The total density of hitchhikers is simply the sum of the two sub-populations discussed above (Supplementary Figure 13). At low and intermediate frequencies the density is dominated by the contribution from *ρ*_1_ i.e. from hitchhikers that occurred on an existing beneficial clone. However at frequencies close to *f* ∼ *ϕ*, the contribution of *ρ*_2_ i.e. hitchhikers that derive from a neutral clone acquiring a subsequent beneficial mutation, becomes significant. The comparison of our theoretical predictions for the densities of hitchhikers agrees well with simulated data (Supplementary Figure 14). The dashed orange line in Figure 1e of the main text is obtained using eqn. 17 and dividing through by *µ*_*n*_ and using the parameters *µ*_*b*_ = 7 *×* 10^−5^ and *s* = 0.13.

**Supplementary Figure 13.**
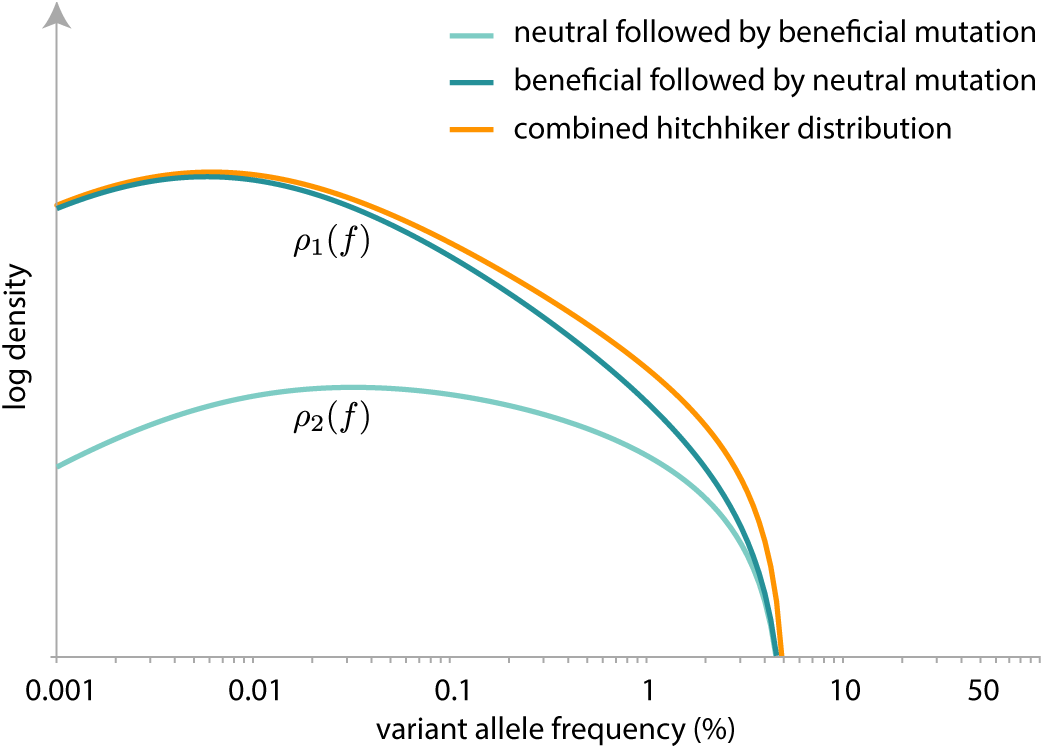
Schematic of the predicted density distribution of hitchhiker variants. The predicted distribution (orange line) is simply the sum of the distributions for the two hitchhiker sub-populations: those in which the beneficial mutation occurred first (dark green line) and those in which the neutral mutation occurred first (light green line).

**Supplementary Figure 14.**
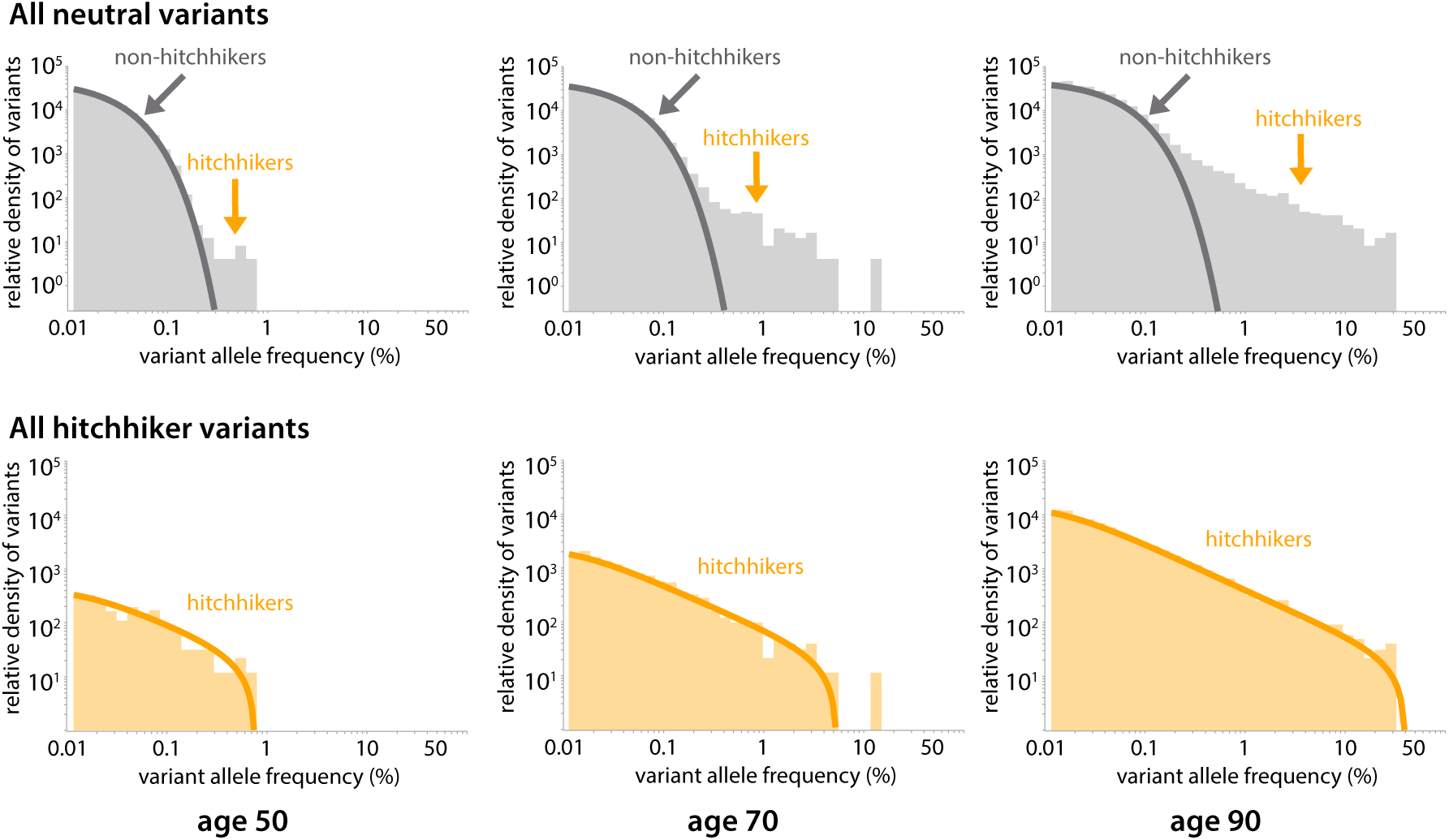
Simulated evolution of HSC clones using HSC total population size of 10^5^, neutral and beneficial mutation rates of 10^−5^ and 10^−6^ per year per cell respectively, symmetric division rate of one per year and fitness effects *s* = 0.1 per year for all beneficial mutations. Simulated hitchhiker densities (orange histogram) results agree closely with theoretical predictions (orange lines) across a range of hitchhiker mutation frequencies and ages.

## Supplementary Note 8: Age prevalence of R882H and R882C mutations

Based on the model, we expect the prevalence of mutations to increase approximately linearly at rate *s × θ*, once the individual is above a certain age determined by the VAF limit of detection (*f*_0_) and the fitness effect (*s*) of the mutation. The reason for this is that, provided *f*_0_ ≪ *ϕ*, the integral can be approximated as:

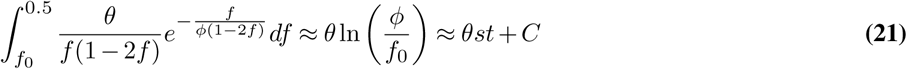

where 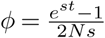 and *C* = *-θ* ln (2*N sf*_0_).

Because the rate of increase in prevalence with age is *s × θ*, examining the age-prevalence relationship of a variant provides us with another method by which the fitness effect, *s* of the variant can be inferred. Two studies (McKerrel 2015^9^ and Coombs 2017^11^) contained sufficient data (*>* 500 total individuals in ≥ 2 age categores) to allow R882 variants to be binned in to sufficient age categories to examine their age-prevalence relationship. R882H and R882C variants were grouped together to increase data strength.

Maximum likelihood estimations were used to calculate the likely fitness effect, *s*, by integrating the expected density of clones between the VAF limit of detection (*f*_0_) and 0.5 across a range of ages, *t*, in Python:

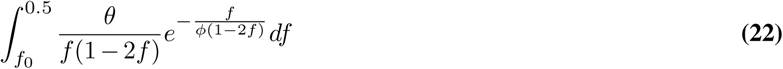

The best-fit *s* for each of the studies were: 12.6% (McKerrel 2015^9^), and 15.7% (Coombs 2017^11^) (Supplementary Figures 15 and 16), which is in good agreement with estimates inferred from the VAF distributions for R882H and R882C (Figure 1e and Supplementary Figure 9). There are very large error bars on these estimates, however, due to the paucity of data in some of the age bins and so these estimates should be treated with caution.

**Supplementary Figure 15.**
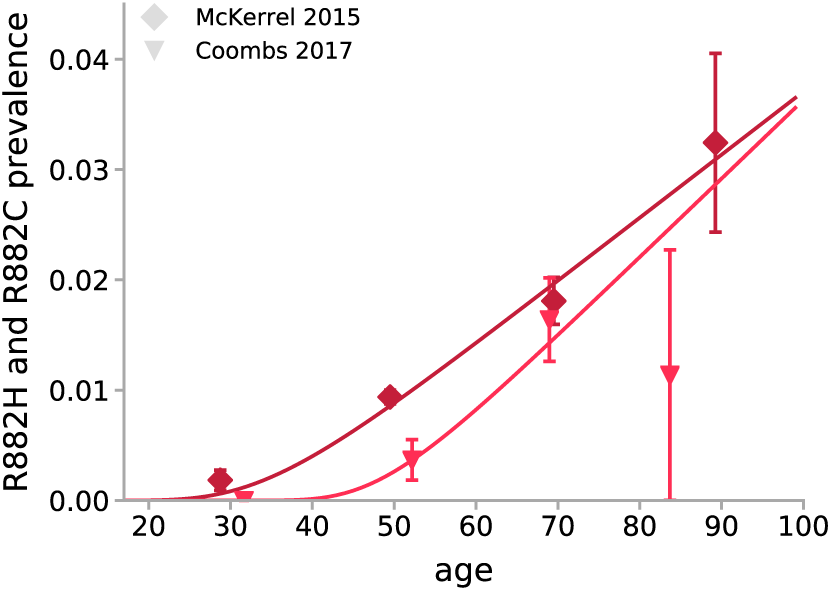
Age-prevalence of R882H and R882C variants. As predicted by the model, the prevalence of R882H and R882C increases approximately linearly with age, once above a certain age determined by the VAF limit of detection and the fitness effect of the mutation. Medium red line: McKerrel 2015. Dark red line: Coombs 2017.

**Supplementary Figure 16.**
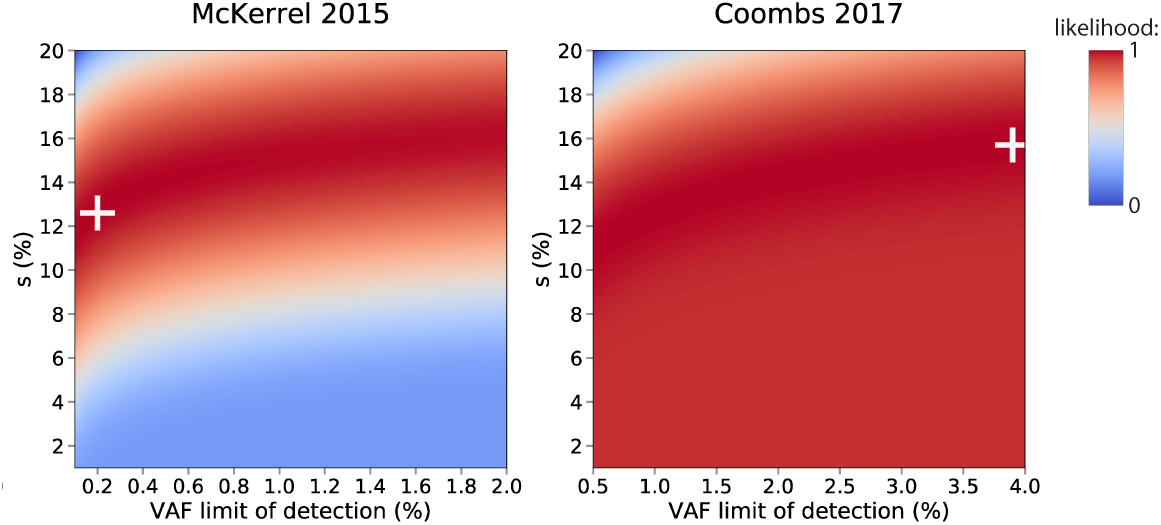
Maximum Likelihood Estimations for fitness effect (*s*) of R882H and R882C mutations from age prevalence data. The most likely fitness effect (*s*) and VAF limit of detection were determined for the two studies that contained sufficient data to analyse their age-prevalence relationship. The white cross marks the most likely *s* and VAF limit of detection for the study.

## Supplementary Note 9: Prevalence of Clonal Haematopoiesis

To estimate the overall prevalence of CH as a function of age, for different sequencing thresholds, we considered the distribution of fitness effects for nonsynonymous variants across 10 of the most commonly mutated CH genes. We considered the regions in these genes that are targeted by a typical sequencing panel, such as the Illumina TruSight Myeloid Panel:

- DNMT3A: all coding exons
- TET2: all coding exons
- ASXL1: exon 12
- JAK2: exon 12 and 14
- TP53: all coding exons
- SF3B1: exons 13 - 16
- SRSF2: exon 1
- IDH2: exon 4
- KRAS: exons 2 - 3
- CBL: exons 8 - 9

The mutation rate, *µ*, for all nonsynonymous variants across these 10 genes was estimated to be 3.77 × 10^−5^, which was calculated by summing the site-specific mutation rates (Table 4) for every possible variant in these regions. Only four studies (Young 2016^10^, Young 2019^15^, Coombs 2017^11^ and Desai 2018^14^) targeted the entirety of these regions and so, when plotting the probability density histogram of variants in these genes, only variants in these four studies were included (Supplementary Figure 17a). As previously, the density of variants were normalised for by dividing by [number of individuals in the study x bin widths] and then rescaled by dividing by 2*µ*.

Estimates for the distribution of fitness effects, *s*, across these 10 genes were inferred by fixing *Nτ* and the standard deviation of ages (*σ*) to that inferred from DNMT3A R882H (Supplementary Note 6). We parameterised the distribution of fitness effects using a family of stretched exponential distributions (eqn. 15) and performed a maximum likelihood procedure, optimising the shape (*β*) and scale (*d*) of the distribution as well as the upper limit *s*_*max*_. This revealed a broad distribution, with 88% of variants having a low fitness and only 2% of variants with a high fitness (Supplementary Figure 17).

The prevalence of CH across these 10 genes, as a function of age, was then calculated using a custom Python script, taking in to account the distribution of fitness effects and different sequencing sequencing sensitivities (VAF limits of detection). Briefly, the mutation rate was normalized by the distribution of fitness effects and, for a given age, the theoretical density (eqn. 9) was integrated over the distribution of fitness effects and over the range of VAFs capable of being sequenced (from the VAF limit of detection to 0.499). The predicted prevalence was then plotted as a function of age for different VAF limits of detection (Figure 3a)

**Supplementary Figure 17.**
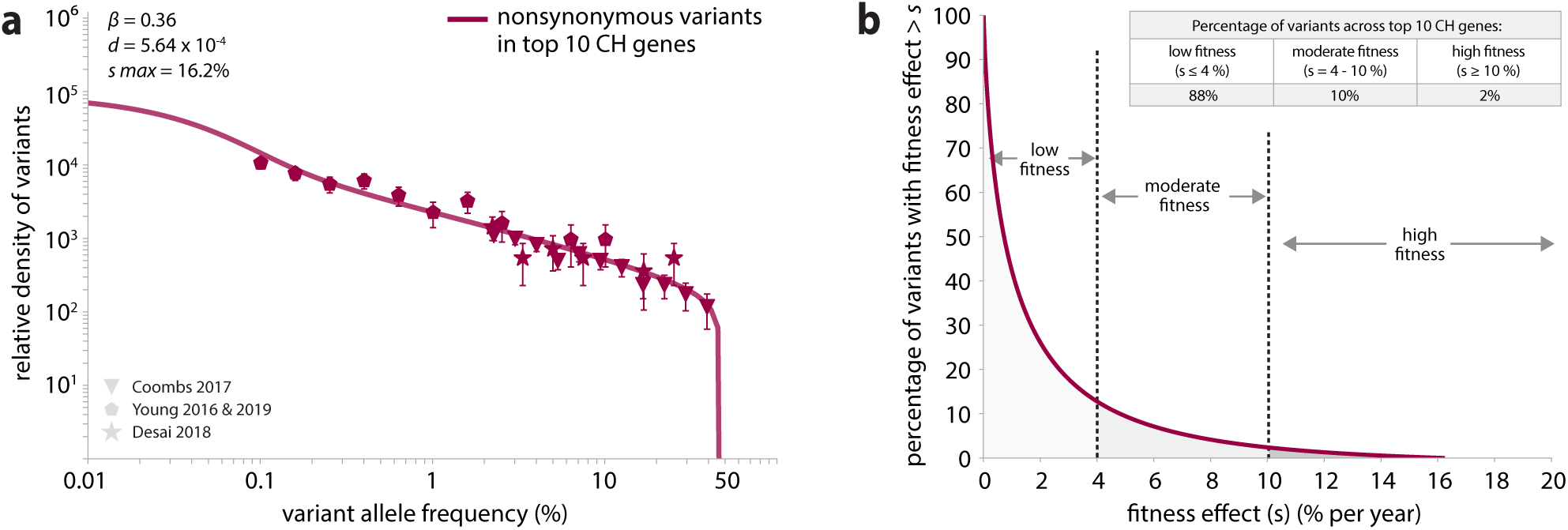
Parameter estimation for the distribution of fitness effects across 10 of the most commonly mutated CH genes: DNMT3A, TET2, ASXL1, JAK2, TP53, SF3B1, SRSF2, IDH2, KRAS, CBL. **(a)**. Probability density histogram for all nonsynonymous variants in these genes. Only four studies (Young 2016^10^, Young 2019^15^, Coombs 2017^11^ and Desai 2018^14^) targeted the entirety of the key coding exons in these genes and so only variants in these four studies were plotted. **(b)** Distribution of fitness effects across these 10 genes.

## Supplementary Note 10: Estimating fitness effects of infrequently mutated sites

In order to determine the fitness effect of individual variants from their VAF-density histograms, our method requires that a variant be seen in at least ~10 individuals. This means that, even with our combined study size of ~ 50,000 individuals, we would have insufficient data to calculate the fitness effects of infrequently mutated variants, even if they were highly fit (Supplementary Note 11). A crude method of determining the potential fitness of variants is to simply determine what fitness effect, given it’s site-specific mutation rate, would be required to explain the number of times the variant is observed.

To crudely determine the fitness effects of variants a cross a gene, e.g. D NMT3A, a custom Python script was used. Briefly, we first created a list of al l the possible nonsynonymous variants within the gene, as well as study-specific lists of variants included in each study’s panel ‘footprint’ of that gene. If a variant was included in a study’s panel, the number of times the variant was expected to be observed in that study was calculated, taking in to account the study’s VAF limit of detection, study size, distribution of ages and the variant’s site-specific mutation rate. This involved integrating the theoretical density (eqn. 9) over the range of VAFs capable of being sequenced by the study (from the VAF limit of detection to 0.499) and then integrating over the distribution of ages for that study. The expected number of observations of that variant was then summed across all the studies which included it in their panel and this number was compared to the actual number of times the variant was observed across all studies. A maximum likelihood approach was then used to determine what fitness effect minimized the L2 norm between the expected and actual number of observations of each variant.

We used this crude counting method to determine the fitness effects of variants Seen more than twice, across all nine studies, in the genes DNMT3A, TET2, ASXL1 and TP53. A limitation of this method is that it does not allow for deviations from the site-specific mutation rates estimated from trinucleotide context, due to its inability to separate out the effects of mutation rate and fitness (in contrast to our VAF-density histogram-based method). Notwithstanding the potential effect this could have on the fitness effect inferred, the counting method suggests there are a number of sites within these four commonly mutated genes that are highly fit yet infrequently mutated (Supplementary Figures 18-21 and Tables 7-10).

**Supplementary Figure 18.**
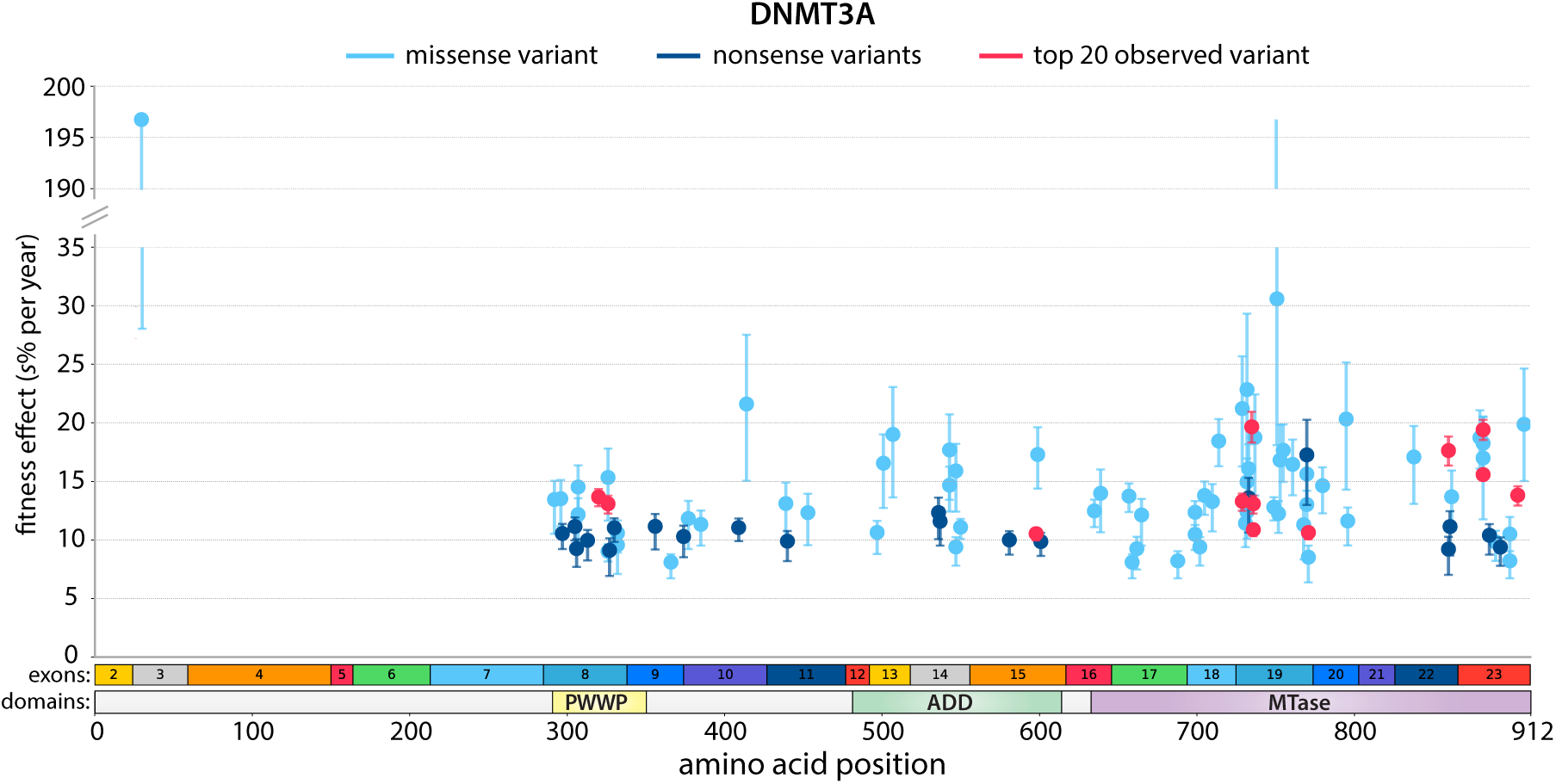
Distribution of fitness effects across DNMT3A, estimated using a crude counting method to infer the fitness effect required to achieve the actual number of observations of the variant. Variants that are in the list of the top 20 most commonly observed variants in CH (from Figure 3) are highlighted in red. Fitness effects were calculated only for those variants observed more than twice across all nine studies.

**Supplementary Figure 19.**
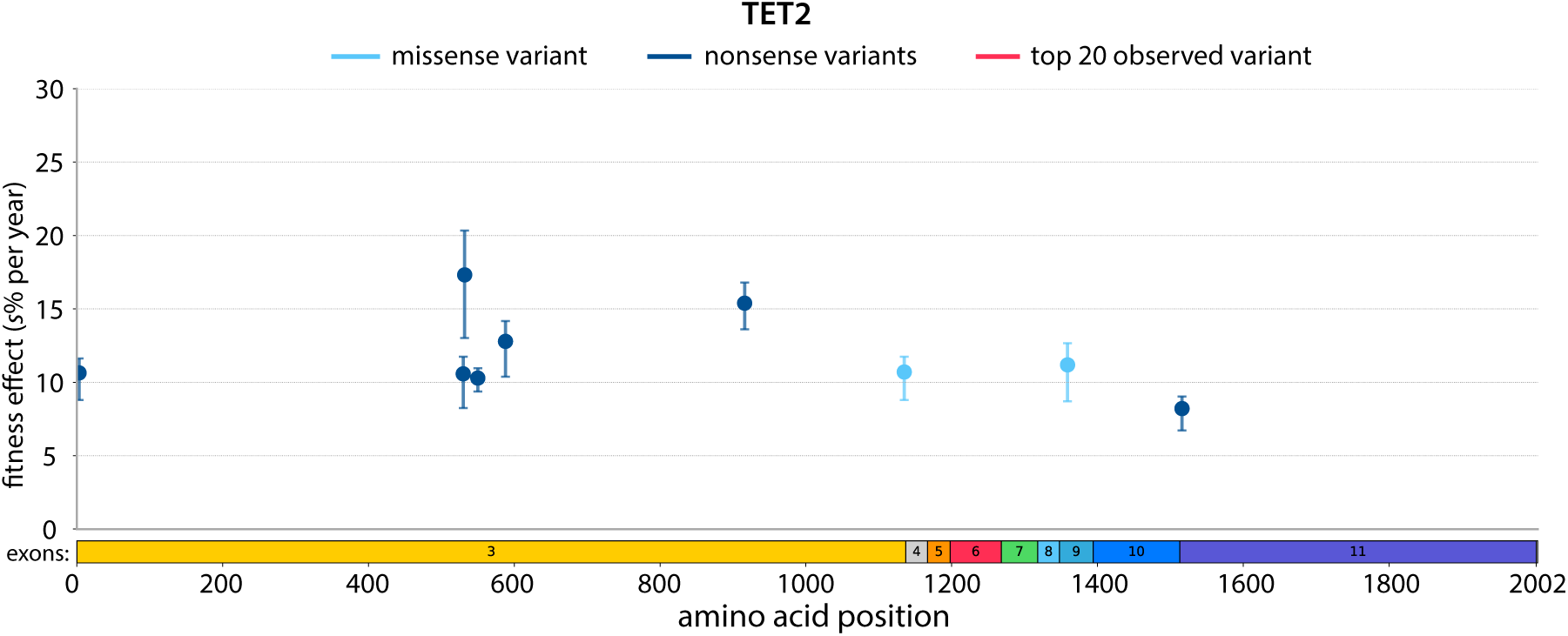
Distribution of fitness effects across TET2, estimated using a crude counting method to infer the fitness effect required to achieve the actual number of observations of the variant. No variants in TET2 were in the top 20 most commonly observed in CH. Fitness effects were calculated only for those variants observed more than twice across all nine studies.

**Supplementary Figure 20.**
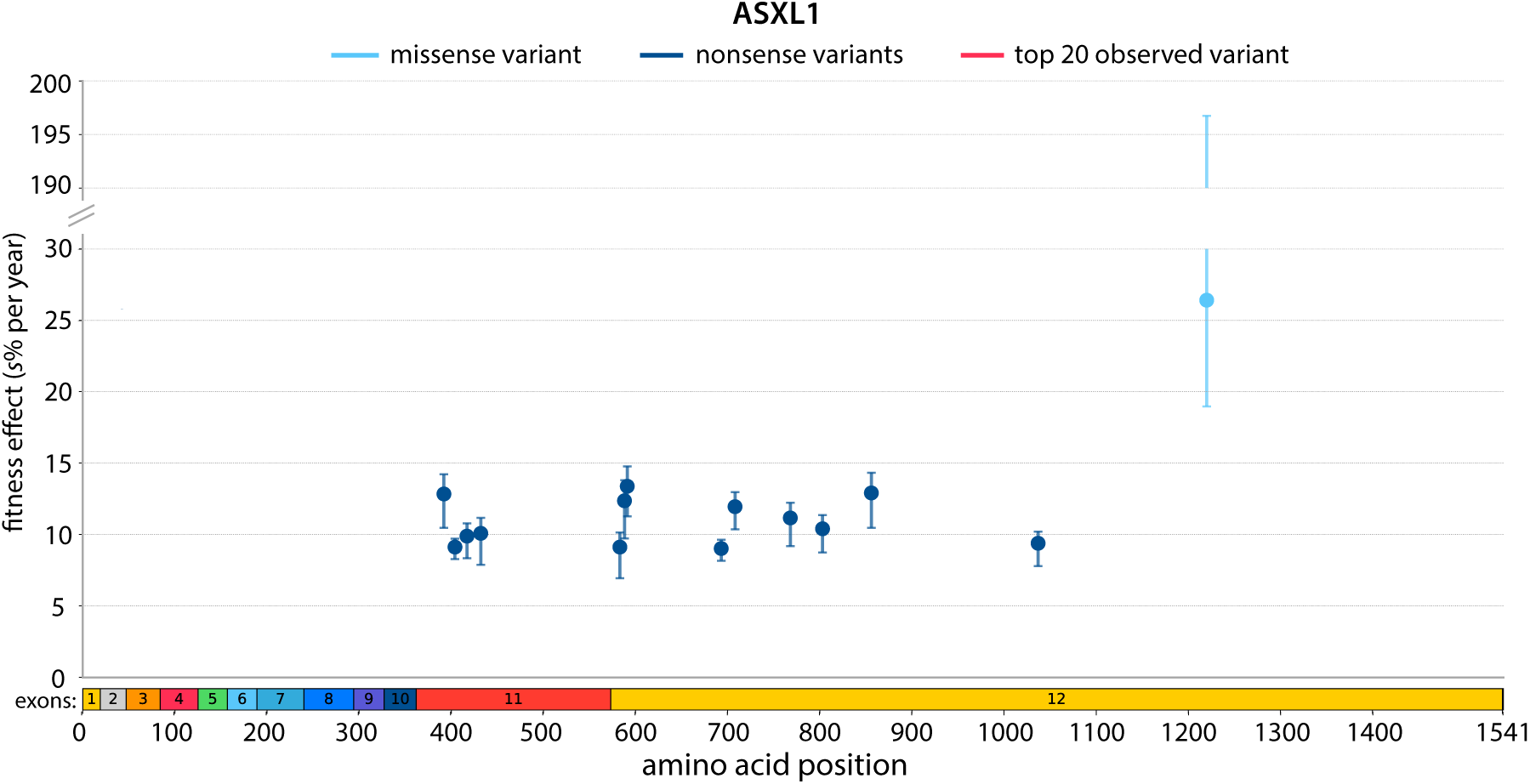
Distribution of fitness effects across ASXL1, estimated using a crude counting method to infer the fitness effect required to achieve the actual number of observations of the variant. No variants in ASXL1 were in the top 20 most commonly observed in CH. Fitness effects were calculated only for those variants observed more than twice across all nine studies.

**Supplementary Figure 21.**
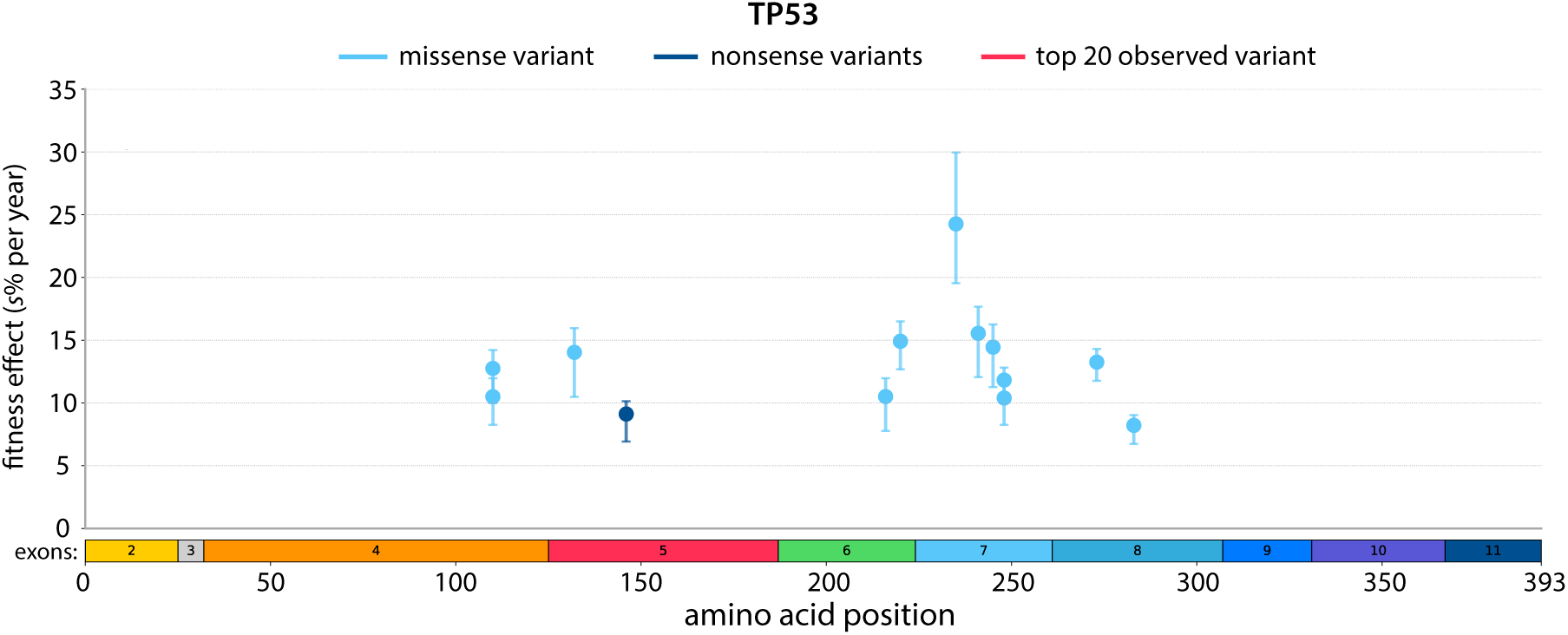
Distribution of fitness effects across TP53, estimated using a crude counting method to infer the fitness effect required to achieve the actual number of observations of the variant. No variants in TP53 were in the top 20 most commonly observed in CH. Fitness effects were calculated only for those variants observed more than twice across all nine studies.

**Table 7.**
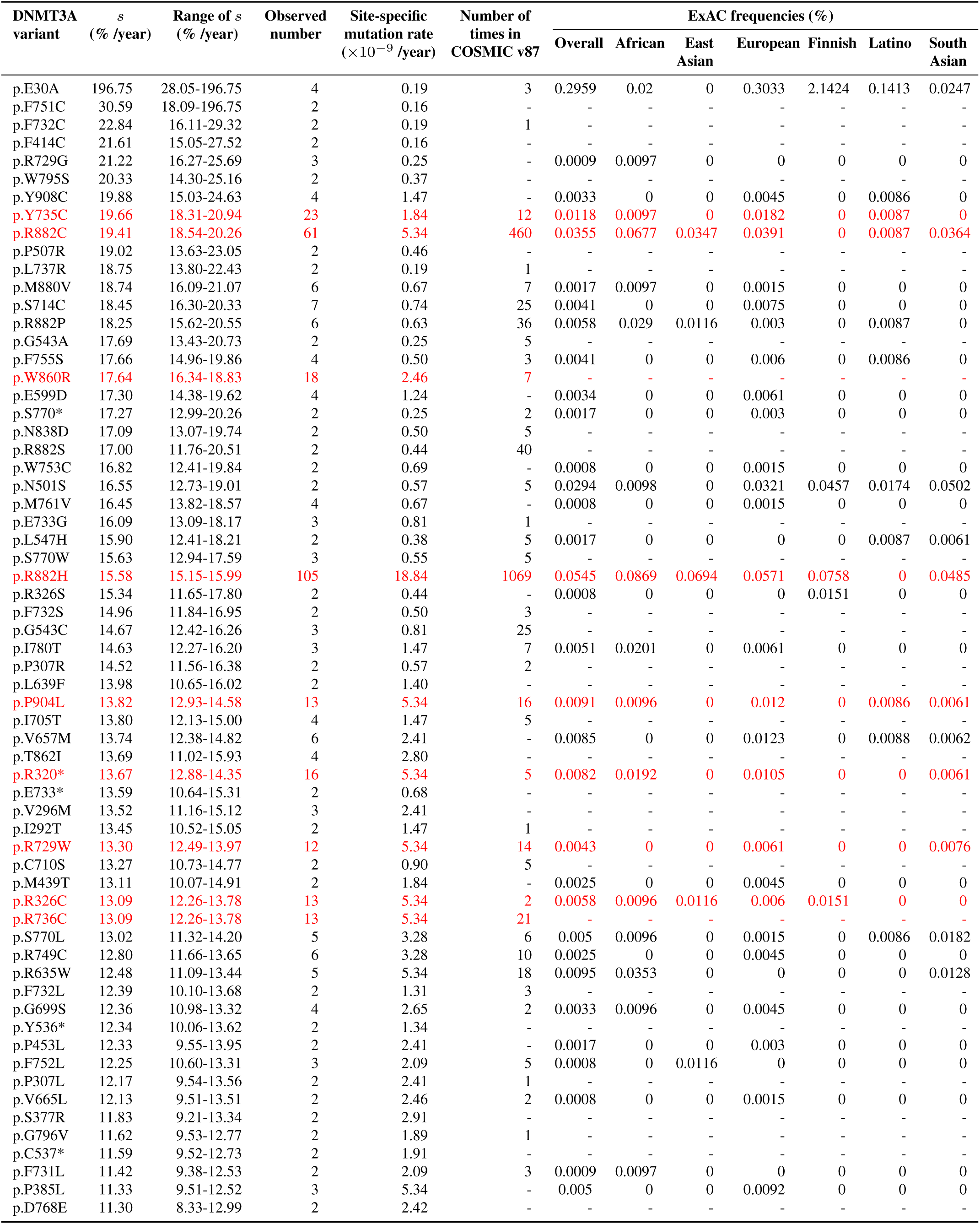
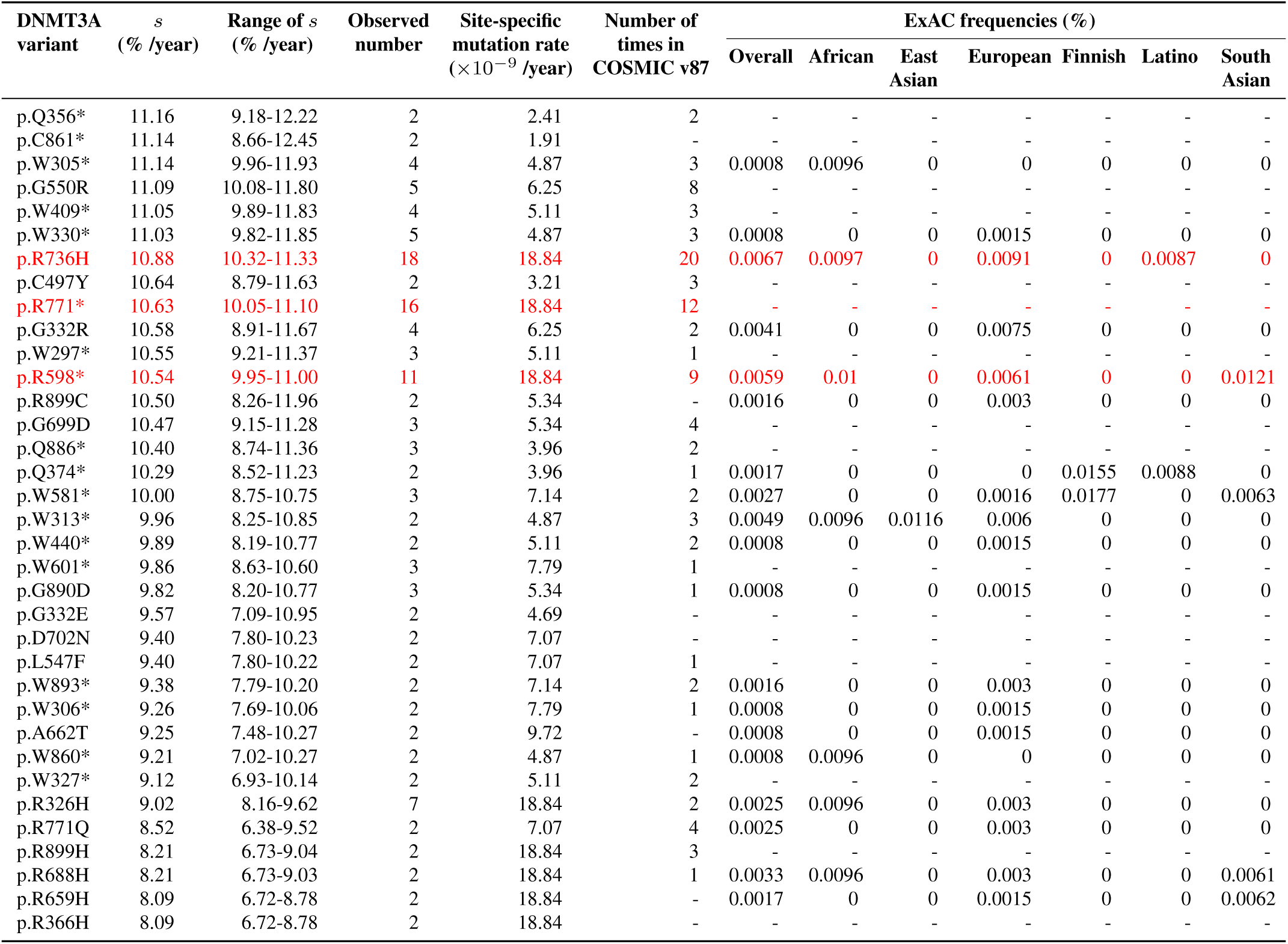
Fitness effects of DNMT3A variants estimated using a crude counting method to infer the fitness effect required to achieve the actual number of observations of the variant. Variants that are within the top 20 observed variants in CH are highlighted in red. Range of *s* was calculated using the sampling noise of the number of observed variants. Site-specific mutation rates are those calculated from trinucleotide context (Supplementary Table 4). The number of times the variant is seen in COSMIC v87 ^60^ (haematopoietic and lymphoid cancers) as well as their frequencies in ExAC are shown.

**Table 8.** Fitness effects of TET2 variants estimated using a crude counting method to infer the fitness effect required to achieve the actual number of observations of the variant. Range of *s* was calculated using the sampling noise of the number of observed variants. Site-specific mutation rates are those calculated from trinucleotide context (Supplementary Table 4). The number of times the variant is seen in COSMIC v87 ^60^ (haematopoietic and lymphoid cancers) as well as their frequencies in ExAC are shown.

| TET2 variant | *s* (% /year) | Range of *s* (% /year) | Observed number | Site-specific mutation rate ($\times 10^{-9}$ /year) | Number of times in COSMIC v87 | ExAC frequencies (%) | | | | | | |
| --- | --- | --- | --- | --- | --- | --- | --- | --- | --- | --- | --- | --- |
|  |  |  |  |  |  | Overall | African | East Asian | European | Finnish | Latino | South Asian |
| p.L532* | 17.32 | 13.02 - 20.34 | 2 | 0.25 | - | - | - | - | - | - | - | - |
| p.Q916* | 15.39 | 13.61 - 16.80 | 5 | 1.09 | 43 | 0.0008 | 0 | 0 | 0 | 0 | 0 | 0.0061 |
| p.S588* | 12.79 | 10.38 - 14.17 | 2 | 1.10 | 4 | - | - | - | - | - | - | - |
| p.R1359H | 11.19 | 8.70 - 12.66 | 2 | 3.96 | 6 | - | - | - | - | - | - | - |
| p.C1135Y | 10.70 | 8.79 - 11.75 | 2 | 3.21 | 3 | 0.0008 | 0 | 0 | 0 | 0 | 0.0087 | 0 |
| p.Q3* | 10.64 | 8.79 - 11.63 | 2 | 3.21 | - | - | - | - | - | - | - | - |
| p.Q530* | 10.58 | 8.24 - 11.74 | 2 | 2.41 | 4 | 0.0008 | 0 | 0 | 0.0015 | 0 | 0 | 0 |
| p.R550* | 10.29 | 9.37 - 10.96 | 7 | 9.72 | 61 | 0.0058 | 0 | 0.0116 | 0.0075 | 0 | 0.0087 | 0 |
| p.R1516* | 8.21 | 6.73 - 9.04 | 2 | 18.84 | 26 | - | - | - | - | - | - | - |

**Table 9.** Fitness effects of ASXL1 variants estimated using a crude counting method to infer the fitness effect required to achieve the actual number of observations of the variant. Range of *s* was calculated using the sampling noise of the number of observed variants. Site-specific mutation rates are those calculated from trinucleotide context (Supplementary Table 4). The number of times the variant is seen in COSMIC v87 ^60^ (haematopoietic and lymphoid cancers) as well as their frequencies in ExAC are shown.

| ASXL1 variant | $s$<br>(% /year) | Range of $s$<br>(% /year) | Observed number | Site-specific mutation rate<br>($\times 10^{-9}$ /year) | Number of times in COSMIC v87 | ExAC frequencies (%) | | | | | | |
| --- | --- | --- | --- | --- | --- | --- | --- | --- | --- | --- | --- | --- |
|  |  |  |  |  |  | Overall | African | East Asian | European | Finnish | Latino | South Asian |
| p.I1220F | 26.41 | 18.97 - 196.75 | 3 | 0.31 | 1 | 0.0099 | 0 | 0 | 0.0181 | 0 | 0 | 0 |
| p.Y591* | 13.38 | 11.27 - 14.77 | 3 | 1.11 | 23 | 0.0025 | 0 | 0 | 0.0046 | 0 | 0 | 0 |
| p.C856* | 12.91 | 10.47 - 14.32 | 2 | 1.04 | 1 | 0.0008 | 0 | 0 | 0.0015 | 0 | 0 | 0 |
| p.S392* | 12.84 | 10.46 - 14.21 | 2 | 1.10 | - | - | - | - | - | - | - | - |
| p.Q588* | 12.36 | 9.72 - 13.80 | 2 | 1.09 | 11 | - | - | - | - | - | - | - |
| p.Q708* | 11.95 | 10.36 - 12.96 | 3 | 2.41 | 8 | - | - | - | - | - | - | - |
| p.Q768* | 11.16 | 9.18 - 12.22 | 2 | 2.41 | 3 | 0.0017 | 0 | 0 | 0.003 | 0 | 0 | 0 |
| p.Q803* | 10.40 | 8.74 - 11.36 | 3 | 3.96 | 2 | 0.0025 | 0 | 0 | 0.0015 | 0 | 0.0174 | 0 |
| p.Q432* | 10.08 | 7.88 - 11.16 | 2 | 3.21 | - | 0.0025 | 0 | 0 | 0.0045 | 0 | 0 | 0 |
| p.R417* | 9.88 | 8.33 - 10.78 | 3 | 5.34 | 8 | 0.0074 | 0 | 0.0116 | 0.009 | 0.0302 | 0 | 0 |
| p.W1037* | 9.38 | 7.79 - 10.20 | 2 | 7.14 | 1 | - | - | - | - | - | - | - |
| p.R404* | 9.12 | 8.28 - 9.71 | 7 | 18.84 | 6 | 0.0058 | 0.0096 | 0.0116 | 0.006 | 0.0151 | 0 | 0 |
| p.W583* | 9.12 | 6.93 - 10.14 | 2 | 5.11 | 3 | 0.0017 | 0 | 0 | 0.003 | 0 | 0 | 0 |
| p.R693* | 9.02 | 8.16 - 9.62 | 7 | 18.84 | 75 | 0.005 | 0.0098 | 0.0233 | 0.0046 | 0 | 0 | 0 |

**Table 10.** Fitness effects of TP53 variants estimated using a crude counting method to infer the fitness effect required to achieve the actual number of observations of the variant. Range of *s* was calculated using the sampling noise of the number of observed variants. Site-specific mutation rates are those calculated from trinucleotide context (Supplementary Table 4). The number of times the variant is seen in COSMIC v87 ^60^ (haematopoietic and lymphoid cancers) as well as their frequencies in ExAC are shown.

| TP53 variant | $s$<br>(% /year) | Range of $s$<br>(% /year) | Observed number | Site-specific mutation rate<br>($\times 10^{-9}$ /year) | Number of times in COSMIC v87 | ExAC frequencies (%) | | | | | | |
| --- | --- | --- | --- | --- | --- | --- | --- | --- | --- | --- | --- | --- |
|  |  |  |  |  |  | Overall | African | East Asian | European | Finnish | Latino | South Asian |
| p.N235S | 24.27 | 19.53 - 29.95 | 5 | 0.62 | 6 | 0.0239 | 0 | 0 | 0.0345 | 0.0913 | 0 | 0 |
| p.S241C | 15.55 | 12.05 - 17.67 | 2 | 0.74 | 9 | 0.0008 | 0 | 0 | 0 | 0.0152 | 0 | 0 |
| p.Y220C | 14.91 | 12.67 - 16.50 | 4 | 1.47 | 145 | 0.0025 | 0 | 0 | 0.0045 | 0 | 0 | 0 |
| p.G245C | 14.45 | 11.27 - 16.26 | 2 | 1.04 | 10 | - | - | - | - | - | - | - |
| p.K132E | 14.03 | 10.48 - 15.97 | 2 | 0.95 | 9 | 0.0008 | 0 | 0 | 0.0015 | 0 | 0 | 0 |
| p.R273H | 13.25 | 11.77 - 14.30 | 5 | 3.96 | 106 | 0.0026 | 0 | 0 | 0.0047 | 0 | 0 | 0 |
| p.R110L | 12.75 | 9.98 - 14.22 | 2 | 1.91 | 7 | 0.0017 | 0 | 0 | 0.0015 | 0 | 0 | 0.0061 |
| p.R248Q | 11.82 | 10.29 - 12.82 | 4 | 5.60 | 189 | 0.0058 | 0 | 0.0231 | 0.0075 | 0 | 0 | 0 |
| p.V216M | 10.51 | 7.77 - 11.96 | 2 | 3.21 | 60 | - | - | - | - | - | - | - |
| p.R110C | 10.50 | 8.26 - 11.96 | 2 | 5.34 | - | 0.0008 | 0 | 0 | 0 | 0.0151 | 0 | 0 |
| p.R248W | 10.39 | 8.26 - 11.56 | 2 | 5.34 | 129 | 0.0008 | 0 | 0 | 0.0015 | 0 | 0 | 0 |
| p.W146* | 9.12 | 6.93 - 10.14 | 2 | 5.11 | 9 | - | - | - | - | - | - | - |
| p.R283H | 8.21 | 6.73 - 9.04 | 2 | 18.84 | 2 | 0.0066 | 0 | 0 | 0.003 | 0 | 0.0262 | 0.0183 |

## Supplementary Note 11: Limitations of study size and sequencing limit

To determine the fitness effect of a variant from its VAF-density distribution, the variant needs to be seen in at least ∼ 10 individuals. The number of individuals a variant is seen in is determined by the fitness effect of the variant (*s*), its mutation rate (*u*) and the VAF limit detection of the study. To calculate the study size needed to determine the fitness of variants of given fitness effects and mutation rates, as a function of sequencing VAF limit of detection, a custom Python script was used. Briefly, the predicted prevalence, for variants of a given *s* and *µ*, were calculated by integrating the theoretical density (eqn. 9) over the distribution of ages (mean age 55, normally distributed with standard deviation 11.4 years, inferred from R882H variants (Supplementary Note 6)) and then integrated over the range of VAFs capable of being sequenced (from the VAF limit of detection to 0.499). The prevalence was then used to calculate how many individuals would be required in order to observe the variant in at least 10 individuals (= 10/prevalence).

The average site-specific mutation rate is ∼ 1.3 *×* 10^−9^ / year, but site-specific mutation rates range from as low as ∼ 10^−10^ to as high as ∼ 10^−8^ / year (Supplementary Note 4). To determine the fitness of all variants with a selective advantage large enough to expand significantly over a human lifespan (*s >* 4%), even those with very low mutation rates (∼ 10^−10^ / year), a study size of >10,000,000 would be required using standard sequencing methods with a VAF limit of detection of 1% (Supplementary Figure 22a). With more sensitive sequencing (VAF limit 0.01%), a study size of ∼ 300, 000 would be needed. The higher the mutation rate of the variant, the smaller the study is required (Supplementary Figures 22b and c).

**Supplementary Figure 22.**
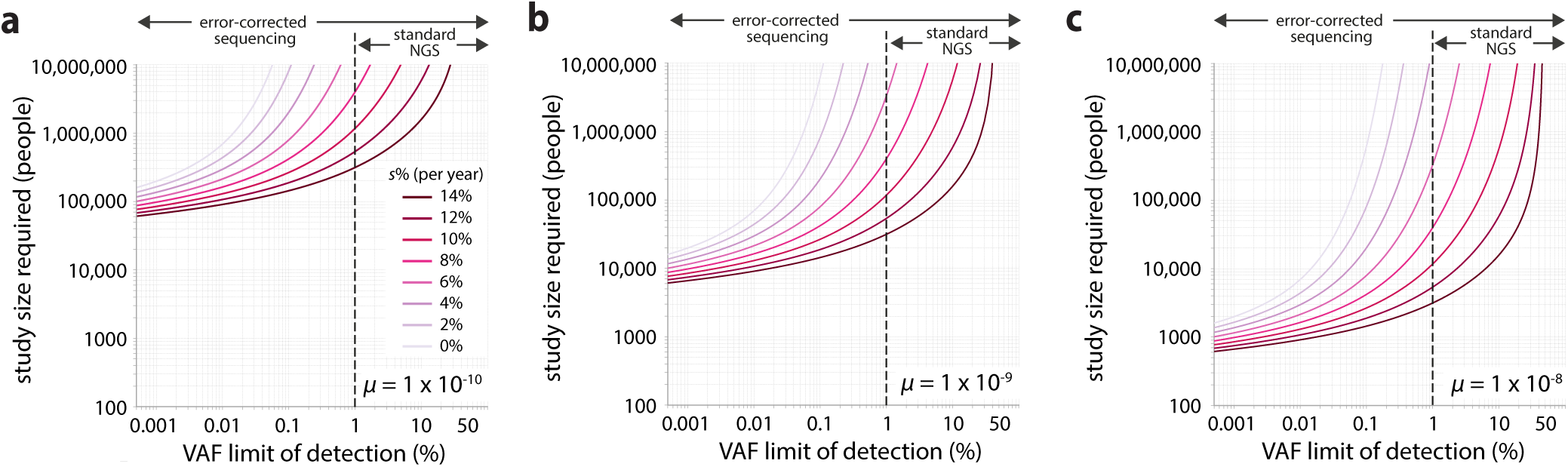
Study size required to accurately quantify different fitness effects (coloured lines) for individual variants, as a function of sequencing sensitivity (VAF limit of detection). **(a)** Study size required for variants with mutation rates of 1 *×* 10^−10^ per year. **(b)** Study size required for variants with mutation rates of 1 *×* 10^−9^ per year. **(c)** Study size required for variants with mutation rates of 1 *×* 10^−8^ per year.

